# Clustering of the Adhesion-GPCR Latrophilin-3 Drives Excitatory Synapse Assembly

**DOI:** 10.64898/2026.10.12.757884

**Authors:** Chuqiao Huyan, Abrar Bhat, Gemma Landa, Thomas C. Südhof, Alexander R. Dunn

**Affiliations:** Department of Chemical Engineering, Stanford University, Palo Alto, CA, USA; Department of Molecular and Cellular Physiology, Stanford University School of Medicine, Stanford, CA 94305, USA; Howard Hughes Medical Institute, Stanford University School of Medicine, Stanford, CA 94305, USA

## Abstract

Precise control of synapse formation is essential for brain function. Adhesion-GPCRs combine adhesion, signaling, and scaffold recruitment to regulate synapse formation, yet their mechanisms of action remain unclear. Here, we identify nanoscale clustering of the adhesion-GPCR Latrophilin-3 as a key mechanism regulating excitatory synapse assembly. We find that endogenous Latrophilin-3 forms densely packed nanoclusters, with larger nanoclusters enriched at excitatory synapses. To probe the functional significance of Latrophilin-3 clustering, we engineered an optogenetically controlled Latrophilin-3. Light-induced Latrophilin-3 clustering triggered robust excitatory synapse formation, which required both Latrophilin-3-mediated G-protein signaling and postsynaptic scaffold recruitment. Strikingly, clustering of a mutant Latrophilin-3 lacking known ligand-binding domains still promoted synapse formation, indicating that Latrophilin-3 ligands induce synapse assembly primarily by clustering Latrophilin-3. Our findings suggest a general paradigm by which extracellular ligand-induced clustering activates adhesion-GPCRs by generating compartmentalized signaling, which in the case of Latrophilin-3, cooperates with postsynaptic scaffold recruitment to drive synapse assembly.

## Introduction

Synapse formation underlies all brain functions, from establishing the initial circuit architecture of the brain to the continuous remodeling of this architecture throughout life. Synaptic adhesion molecules (SAMs) have emerged as key organizers for synapse formation^1,2^. Accumulating evidence suggests that particular SAMs specify where and when synapses form and determine the functional properties of synapses^3,4^. Although enormous progress in characterizing diverse SAMs has been achieved in recent years, the general principles governing synapse assembly and synapse specification remain incompletely understood.

An emerging view is that synapse assembly depends not only on the molecular composition of synapses, but also on the spatial organization of synaptic proteins. Recent super-resolution studies have revealed that presynaptic active zone proteins and postsynaptic scaffolding proteins form nanoclusters that are aligned in nanocolumns, whose size and number may modulate synaptic transmission and plasticity^5–8^. In a prominent example, it was shown that presynaptic neurexins and their postsynaptic LRRTM and neuroligin ligands are organized into nanoclusters at synapses and that these SAMs control the localization of neurotransmitter receptors^9–13^, although the precise relation of the nanoclusters to the neurotransmitter receptor organization has remained unclear. Thus, comparatively little is known about how spatial arrangement of SAMs may generally regulate synapse assembly.

Here, we address this question by focusing on Latrophilin-3 (Lphn3; gene symbol *ADGRL3*), an adhesion-G protein–coupled receptor (adhesion-GPCR) associated with psychiatric disorders such as ADHD and substance use disorder^14–17^. Lphn3 is a key actor in excitatory synapse formation whose expression and alternative splicing are developmentally controlled, activity-dependent, and regulated in a cell-type specific manner^18–20^. Previous studies have shown that the function of Lphn3 in synapse assembly relies on its ability to i) bind extracellular ligands (specifically teneurins and fibronectin-leucine-rich transmembrane proteins [FLRTs]), ii) to activate Gα_S_-dependent GPCR signaling, and iii) to interact with postsynaptic scaffolding proteins such as SHANKs via its intracellular PDZ-binding motif (PBM)^19–21^. These studies established that Lphn3 is a key multifunctional synapse organizer, but how ligand binding controls Lphn3 function and how intracellular G-protein signaling and scaffolding protein recruitment mediated by Lphn3 promote synapse formation is unknown.

In the present study we find that, using super-resolution microscopy, Lphn3 assembles into densely packed nanoclusters. Larger clusters preferentially localize near or within excitatory synapses and correlate with synaptic transmission strength. To test whether Lphn3 clustering plays a causal role in synapse formation, we engineered an optogenetic Lphn3 construct that enables acute, light-induced Lphn3 nanoclustering. Induced clustering of exogenous Lphn3 robustly promoted excitatory synapse formation and enhanced network transmission in cultured neurons even in the presence of endogenous Lphn3. Mechanistically, clustering-induced synapse assembly required Lphn3-mediated activation of heterotrimeric G proteins and engagement of postsynaptic density scaffolds via its PDZ-binding motif but bypassed the need for extracellular ligand binding domains that are otherwise essential for Lphn3-dependent synapse formation. Together, these results identify Lphn3 nanoscale clustering as a key mechanism triggering synapse formation, suggesting that coincident nanoscale clustering of postsynaptic scaffolds and heterotrimeric G protein signaling is sufficient to drive excitatory synapse assembly. More broadly, our work suggests a general model for how adhesion-GPCRs may couple extracellular ligand-binding to spatially restricted intracellular signaling and to the recruitment of postsynaptic scaffolds in order to control synapse assembly.

## Results

### Endogenous Lphn3 forms densely packed nanoclusters near excitatory synapses

We first examined the spatial organization of Lphn3 using three-dimensional super-resolution Stimulated Emission Depletion (3D-STED) microscopy. Hippocampal neurons were cultured from Lphn3 conditional KO (cKO) mice in which endogenous Lphn3 is tagged with an N-terminal HA epitope and Lphn3 can be deleted by Cre recombinase^19^ (Fig. 1A). To identify excitatory synapses, we co-stained the neurons for the endogenous Lphn3’s HA epitope, presynaptic vGluT1, and postsynaptic Homer1. STED imaging revealed that Lphn3 assembled into nanoclusters of variable sizes with an average diameter of ∼220 nm (Fig. S1A), whereas synapses as marked by vGluT1 and Homer1 exhibited an average diameter of ∼570 nm and ∼480 nm, respectively (Fig. S1B,C). Notably, Lphn3 clusters were detected not only within excitatory synapses but also adjacent to and distant from excitatory synapses. Sometimes multiple Lphn3 clusters were found associated with a single synaptic junction (Fig. 1B). Importantly, Lphn3 nanoclusters that overlap with synaptic markers are significantly larger in volume (approximately 2-fold increase) than those distant from synaptic markers (Fig. 1B). Thus, larger Lphn3 clusters preferentially localize within or near synapses.

**Figure 1.**
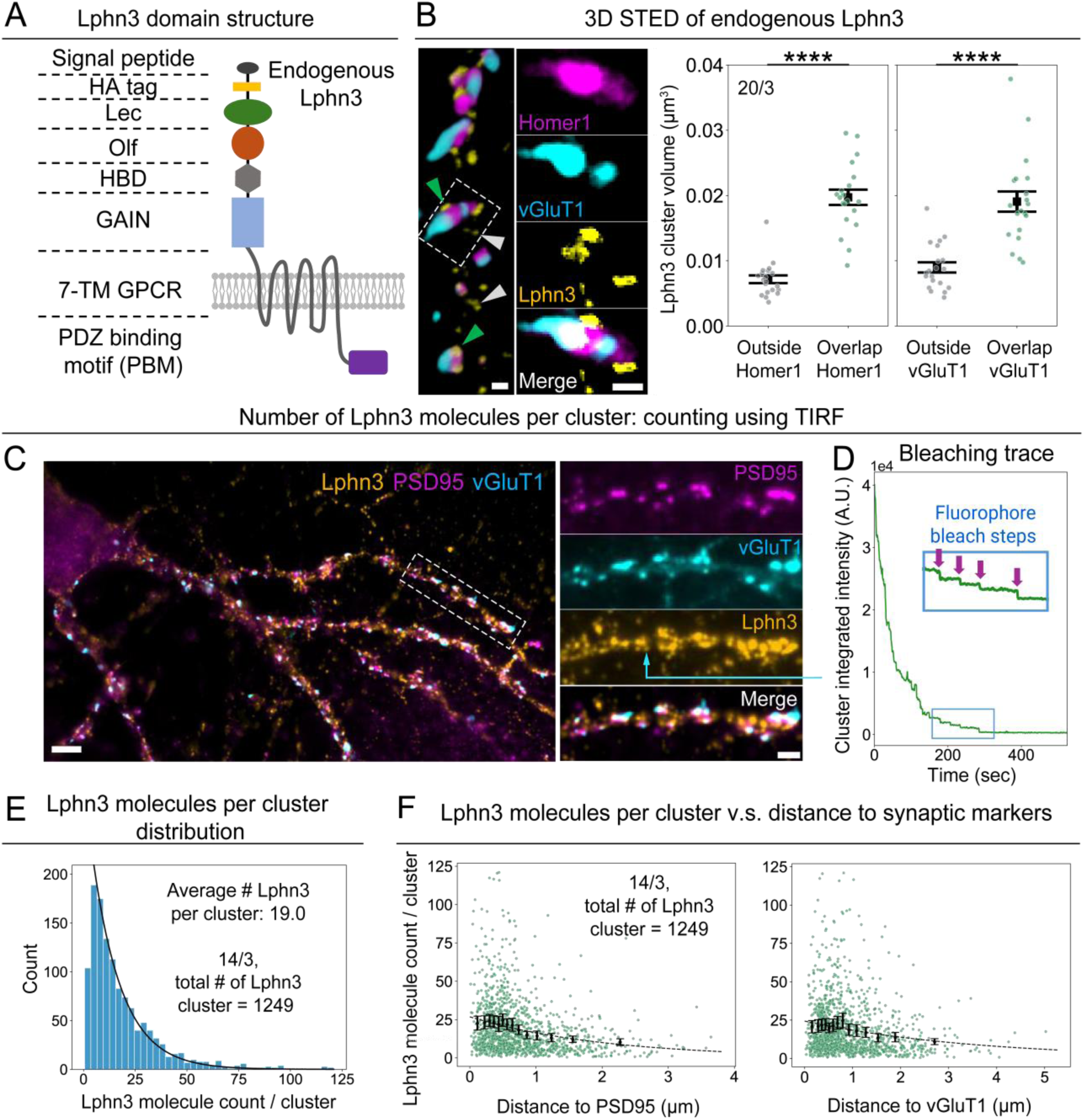
Large Lphn3 nanoclusters are enriched at excitatory synapses. **(A)** Lphn3 domain structure. **(B)** Synaptic Lphn3 nanoclusters are twice as large as non-synaptic Lphn3 nanoclusters on average. *Left.* Example 3D STED image of cultured mouse hippocampal neurons expressing endogenous HA-tagged Lphn3 (green and white arrow heads denote Lphn3 clusters within and outside of excitatory synapses, respectively; raw images show a 3D view (dendrite overview) or one z slice (zoom-ins); scale bars = 0.5 μm. *Right*. Volume of Lphn3 clusters that overlap with or are outside of the areas of the synaptic markers Homer1 and vGluT1; mean ± SEM; dots are data from each dendritic segment (pseudo-replicates); statistics: independent t-test. \*\*\*\**P* < 0.0001. **(C)** Cultured neurons stained for endogenous Lphn3 and pre- and postsynaptic markers imaged by TIRF. Scale bars: 5 μm (overview), 2 μm (dendritic segment). (**D)**. Example photobleaching trace for a Lphn3 cluster (blue arrow in **C**) to illustrate single-molecule bleaching steps. (**E)**. Estimated Lphn3 cluster size distribution, corrected for the number of dye molecules per antibody (black line, exponential fit for cluster sizes ≥5, with a decay constant = 0.07. (**F)**. Plots of the Lphn3 cluster size vs. distance to postsynaptic PSD95 (*left*) and presynaptic vGluT1 markers (*right*). Data are organized into equal-sized bins. Within each bin, black dots show average and error bars show the 95% confidence interval from bootstrap resampling (1000 iterations). Black dashed line is calculated from an exponential fit of the mean, with decay constants of 0.49 (*left*) and 0.28 μm^-1^ (*right*). Note across all figures, squares and error bars represent mean ± SEM unless specified otherwise; number of technical replicates and biological replicates are labeled on the plots.

To estimate the number of Lphn3 molecules per cluster, we used a photobleaching-based quantification approach that employs high-resolution total internal reflection fluorescence (TIRF) microscopy^22–24^ (Fig. 1C). We labeled endogenous Lphn3 with fluorophore-conjugated HA antibodies and acquired photobleaching traces. These traces exhibited discrete single-molecule bleaching steps (Fig. 1D). After calibrating the average number of fluorophores per antibody (Fig. S1F, G), we estimated the number of Lphn3 molecules per cluster based on the total pre-bleaching brightness. This analysis yielded an average of ∼20 Lphn3 molecules per cluster, with an approximately exponential distribution of cluster sizes (Fig. 1E). The estimate of ∼20 Lphn3 molecules per cluster likely represents a lower boundary due to incomplete labeling. Nonetheless, combining the average Lphn3 cluster size (∼20) and typical cluster dimensions (∼200 nm diameter) implies an average intermolecular spacing of ∼40 nm. This spacing is consistent with the anticipated size of the Lphn3/teneurin/FLRT ternary complex^25–30^, indicating that Lphn3 molecules are densely packed within individual clusters.

We next aimed to further examine the observation obtained by STED imaging that Lphn3 nanoclusters are larger close to than outside of synapses (Fig. 1B). We plotted the estimated number of Lphn3 molecules per cluster from TIRF imaging against the distance of each cluster to the nearest excitatory synapse, defined by the centroid of vGluT1 or PSD95 puncta (Fig. 1F). The results showed that, while clusters with small numbers of Lphn3 molecules were found both near and away from synapses, clusters with larger numbers of Lphn3 molecules were predominantly positioned within or adjacent to excitatory synapses. These data further support the conclusion that larger Lphn3 nanoclusters are preferentially found in synaptic junctions.

Are the larger Lphn3 clusters that preferentially localize to synapses actually associated with functional synapses? To address this question, we combined Lphn3 imaging with functional imaging using the glutamate reporter iGluSnFR3^31^. This approach enabled us to simultaneously identify active excitatory synapses by visualizing glutamate release sites and to monitor Lphn3 nanoclusters at these synapses (Fig. 2A). Because the number of Lphn3 molecules per cluster correlates with cluster fluorescence intensity (Fig. S1H), we used the normalized cluster fluorescence intensity as a proxy for the Lphn3 molecular count per cluster. Again, we observed that brighter Lphn3 clusters preferentially localized near glutamate release sites (Fig. 2B).

**Figure 2.**
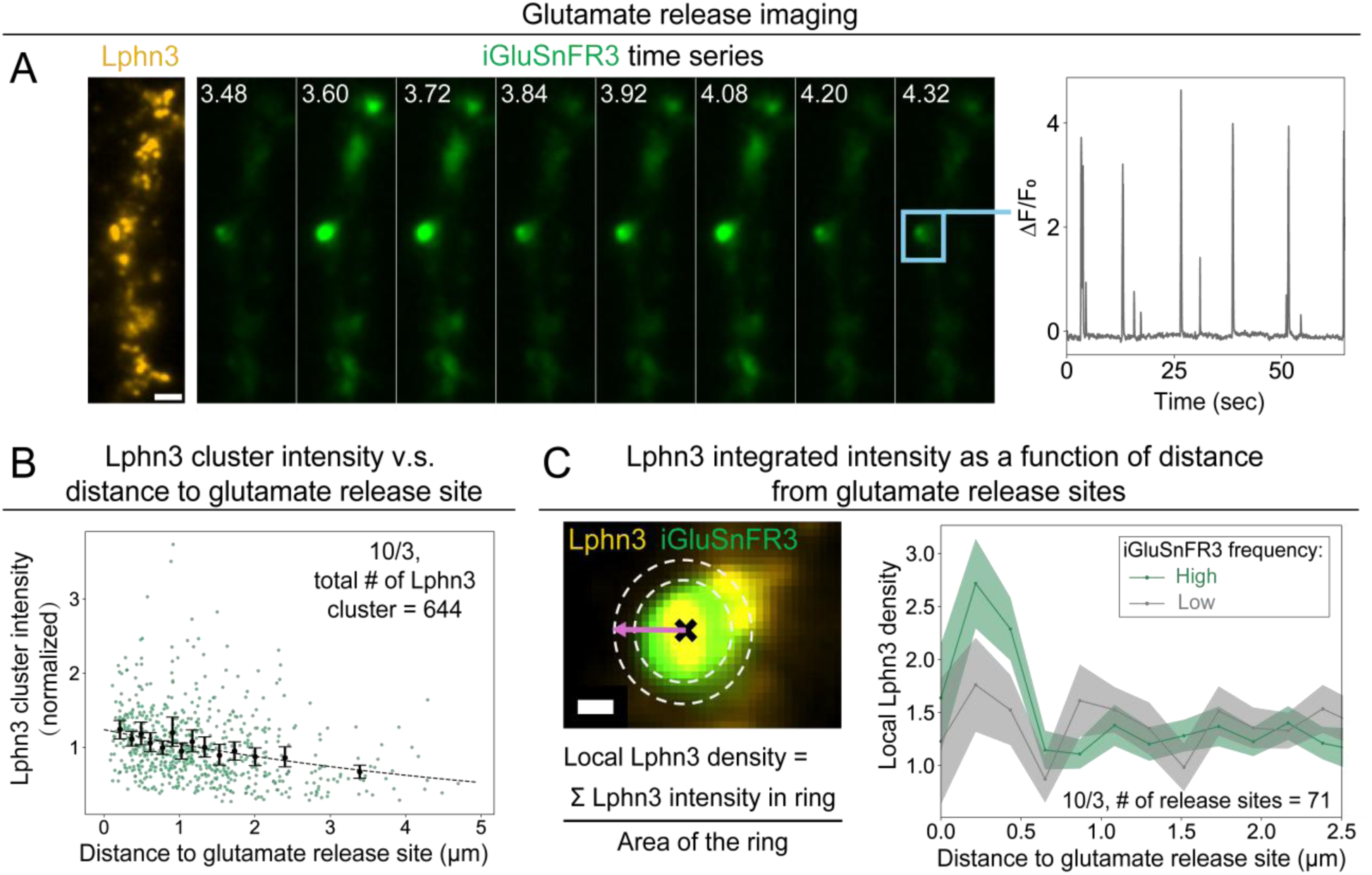
Lphn3 nanocluster size correlates with the activity of glutamate release sites at excitatory synapses. **(A).** Analysis of glutamate release sites using iGluSnFR3 imaging. *Left*. Surface-labeled endogenous Lphn3 (orange) and snapshot images of glutamate imaging using iGluSnFR3 (green). Scale bar: 2 μm. *Right*. Temporal trace of an example glutamate release site (blue square). (**B**). Number of Lphn3 molecules per cluster (estimated by normalized cluster intensity) plotted against the distance to excitatory synapses as defined by glutamate release sites. Data are organized into equal size bins. Within each bin, black dots show average, while error bars show the 95% confidence interval from bootstrap resampling (1000 iterations). Black dashed line is calculated from exponential fit, with a decay constant of 0.17 μm^-1^. **(C)** Calculation of the Lphn3 areal density surrounding glutamate release sites. *Left*. Schematic depicting local Lphn3 density around one glutamate release site as a function of distance (*d*) from the synapse. Scale bar: 0.5 μm. “X” marks the center of a glutamate release site, and the dashed lines denote the ring area that is at a specific distance *d* from the release site (*pink arrow*). *Right*. Local Lphn3 density as function of distance from glutamate release sites. Mean ± SEM is shown. Glutamate release sites are grouped into high (green) and low (grey) glutamate release frequency, with high and low representing the top 75% (high) and bottom 25% (low) of the overall release frequency. In **B** and **C**, Lphn3 intensity and glutamate release frequency are normalized for each dendritic segment to account for variability.

Finally, we asked whether Lphn3 clustering correlates with synaptic strength. Because individual synapses can be associated with multiple Lphn3 clusters, we measured the spatial distribution of Lphn3 around a given glutamate release site using a radial distribution function (RDF), which reports the Lphn3 areal density as a function of distance from synaptic glutamate release sites (Fig. 2C). This analysis revealed an enrichment of Lphn3 near detected glutamate release sites that decayed with distance. Glutamate release sites exhibiting higher activity tend to be associated with a modestly higher local Lphn3 density, defined as within ∼0.6 μm from the release site (Fig. 2C).

Together, these data show that endogenous Lphn3 forms densely packed nanoscale clusters and that larger Lphn3 clusters are preferentially positioned near functional excitatory synapses.

### Optogenetic Lphn3 clustering enhances synapse formation in cultured primary hippocampal neurons

The correlation between the size of Lphn3 clusters and their synaptic vs. non-synaptic localization suggests that Lphn3 clustering at a certain threshold may promote synapse assembly. To directly test this hypothesis, we engineered a Lphn3 construct that enables optogenetic clustering of Lphn3 (Lphn3-Clust). We inserted the CRY2-Clust domain, which oligomerizes upon blue light stimulation^32^, into a relatively non-conserved region of the Lphn3 intracellular sequence (Fig. 3A, Fig. S2A). When Lphn3-Clust was expressed in cultured mouse hippocampal neurons, a single pulse of blue light (2 s) induced the formation of new Lphn3 clusters and the enlargement of pre-existing clusters within 30 seconds (Fig. 3B, Fig. S2B-D, Supplementary Video 1). We confirmed that the Lphn3-Clust protein was functional in synapse formation in the absence of light-induced clustering by demonstrating that Lphn3-Clust rescued the decrease in synapse numbers caused by deletion of endogenous Lphn3 (Fig. 3C, Fig. S3A-B). Lphn3-Clust localized throughout the plasma membrane, with larger clusters primarily found at spine-like structures (Fig. 3B, Fig. 5F), showing a spatial pattern similar to that of endogenous Lphn3.

**Figure 3.**
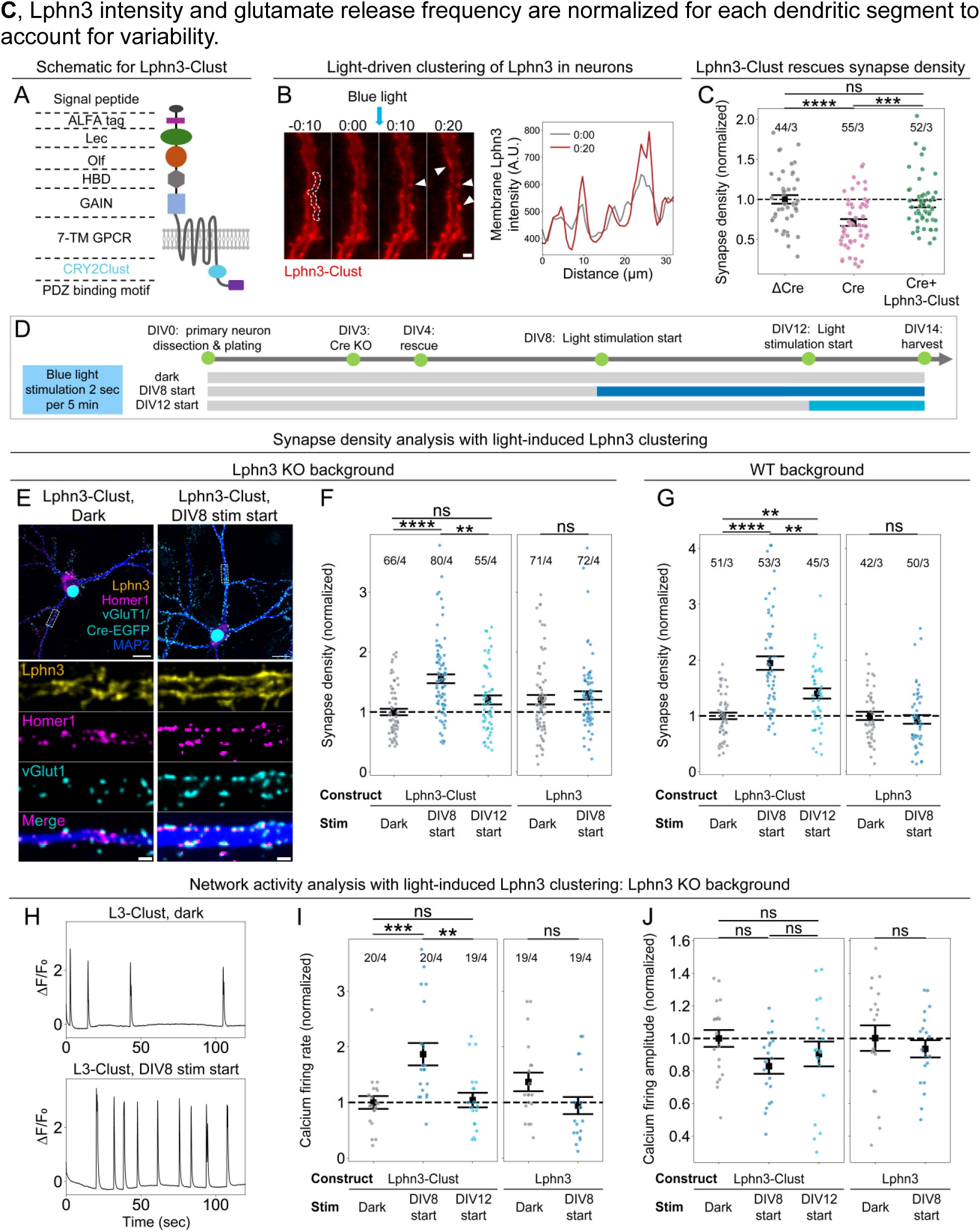
Optogenetic Lphn3 clustering enhances synapse assembly. (**A)** Schematic of the Lphn3-Clust construct in which the CRY2-Clust domain^32^ is inserted into the cytosolic disordered region of Lphn3. (**B)** Light stimulation induces rapid Lphn3-Clust clustering in neurons. *Left.* Representative images (arrow heads: enlarged Lphn3 clusters. Scale bar: 5 μm). *Right.* Lphn3-Clust intensity along a line segment of dendritic membrane, denoted by the white dashed line on the left, before and after light stimulation (line thickness set at 2.59 μm). (**C)** Lphn3-Clust rescues the synapse loss phenotype of Lphn3-deficient neurons without causing by itself an increase in synapse numbers over that of control levels. Excitatory synapse density was measured as overlapping vGluT1 and Homer1 puncta, with Cre (Lphn3 knockout), ΔCre (knockout control), and Cre followed by rescue with Lphn-Clust. (**D)** Timeline of light-induced Lphn3 clustering experiments. **(E-F)** Synapse density analysis with light-induced clustering of Lphn3-Clust in Lphn3 KO neurons. **(E)** Representative images of stained neurons fixed and stained at DIV14. Scale bars: 20 μm (overview)/2 μm (zoom-in). **(F)** Density of excitatory synapses measured by number of puncta positive for both presynaptic vGluT1 and postsynaptic Homer1 markers per μm length of dendrite. **(G)** Synapse density analysis with light-induced clustering of Lphn3-Clust, on WT Lphn3 background. **(H-J) (H)** Representative population calcium imaging traces. (**I-J**). Synchronous firing rate and amplitude quantifications of calcium signals. To account for batch variance, data are normalized to the average value of each batch of the ΔCre condition, Lphn3-Clust dark condition, or dark condition for a specific construct, and this average is denoted by the dashed line. Dots represent data from each dendritic segment (**C, F, G**) or each field-of-view (**I, J**). Statistics: For comparison between more than two conditions, One-way analysis of variance (ANOVA) with post hoc Tukey (equal variance data) or Games-Howell (non-equal variance data) tests were used. For comparison between two conditions, independent t-test was used. \*\*\*\**P* < 0.0001; \*\*\**P* < 0.001; \*\**P* < 0.01; \**P* < 0.05; ns: not significant.

In hippocampal cultures, neurons form synapses beginning at approximately DIV5-6 and continue to assemble synapses over the course of the following 10 days^33^. To apply repeated light stimulation on this timescale and to test different light stimulation conditions in parallel, we engineered a programmable LED device that can apply different light stimulation patterns to separate wells within 24-well plates (Fig. S3C, Supplementary Video 2). Given that the typical disassembly half-life of CRY2-Clust clusters is around 10 minutes^32^, we settled on a protocol in which neurons were exposed to a pulse of blue light every 5 minutes to induce clustering while avoiding light toxicity (Fig. 3D). We applied light stimulation to neurons expressing Lphn3-Clust starting at DIV8 (early development) or at DIV12 (when many synapses had already formed) and used immunocytochemistry and calcium imaging to compare synapse formation under control or light-induced clustering conditions at DIV14. These experiments were performed in neurons lacking endogenous Lphn3 unless noted otherwise (Fig. 3D).

First, we quantified excitatory synapse density by staining for the synaptic markers vGluT1 and Homer1 (Fig. 3E, F). With light-driven clustering applied starting from DIV8, synapse density increased significantly (∼65%) compared to dark conditions. The same light stimulation did not alter synapse density in neurons expressing WT Lphn3, confirming that the observed synapse formation phenotype was not due to light stimulation alone (Fig. 3E, F). We did not detect significant changes in synapse density with light-induced clustering beginning DIV12 (Fig. 3F), suggesting that light-induced Lphn3 clustering is required for synapse formation early on and arguing against a synapse maintenance role for Lphn3. Importantly, when Lphn3-Clust was expressed in the presence of endogenous Lphn3 (i.e., in the absence of the Lphn3 KO), light-induced Lphn3 clustering still robustly increased (∼90%) synapse density (Fig. 3G). This result indicates that Lphn3 clustering can induce synapse assembly even beyond normal levels and can thus act as a dominant-positive agent, which is not observed when Lphn3 is simply overexpressed without clustering^19^.

Receptor clustering can lead to changes in receptor internalization^34,35^, leading us to check whether light-driven Lphn3-Clust clustering changes its surface expression. By measuring surface-labeled Lphn3-Clust receptors in different conditions, we found that light-induced clustering does not significantly alter surface expression of Lphn3-Clust (Fig. S6C).

Second, we asked whether optogenetic Lphn3 clustering results in enhanced neuronal network connectivity, along with increased synapse numbers (Fig. 3H-J). Synchronous calcium spikes measured by imaging in cultured neurons are a proxy for the functional connectivity of neuronal networks^36,37^. Light-induced Lphn3 clustering in neurons lacking endogenous Lphn3 starting at DIV8 led to a significant increase in the synchronous firing rate (∼87%) that was not observed under the dark condition or with light stimulation beginning at DIV12 (Fig. 3H-J). Light stimulation did not alter synchronous firing of neurons expressing WT Lphn3, confirming that the observed phenotype was not due to light stimulation alone (Fig. 3H-J). Light stimulation also led to a small, insignificant decrease in the calcium signal amplitude. These results indicate that increased clustering of Lphn3 during early synapse development enhances both structural and functional maturation of excitatory networks. Note that in these experiments, the non-light condition corresponds to the ‘normal’ state and the increase in synaptic connectivity induced by Lphn3 clustering represents an effective addition of new synapses. The fact that light-induced clustering starting at DIV12 had no effect on synaptic connectivity suggests that the increased number of synapses reflects a Lphn3 function in synapse assembly, not synapse maintenance, and that Lphn3 clustering likely acts by triggering an extended signaling cascade instead of directly stimulating synapse assembly.

### Lphn3 GPCR signaling is required for clustering-induced synapse formation

GPCR signaling by Lphn3 is essential for its role in synapse formation^19,20^. We therefore examined whether Lphn3 GPCR signaling was likewise required for Lphn3 clustering-induced synapse formation. Addressing this question required identification of point mutations in Lphn3 that block heterotrimeric G protein activation while maintaining surface expression levels comparable to those of wild-type (WT) Lphn3. Based on the structure of the Lphn3-Gα_S_ heterotrimer complex^38^ (Fig. 4A, PDB: 7WY8), we introduced into Lphn3 a point mutation (F1041E) that is predicted to impair G protein coupling. Consistent with prior studies^20,39^, TRUPATH signaling assays confirmed that WT Lphn3 activates signaling through Gα_S_, Gα_12_, and Gα_13_, while the F1041E mutant reduces signaling through all three Gα proteins (Fig. 4B). Given that activation of the Gα_S_ pathway is particularly important for Lphn3’s role in synapse assembly^20,33^, we measured the downstream cAMP activity induced by WT-Lphn3 and F1041E-Lphn3 expression in HEK cells. Using the cAMP reporter G-Flamp1^40^, we observed that WT but not F1041E-mutant Lphn3 strongly increased cAMP levels (Fig. 4C). Importantly, surface expression levels of WT and mutant Lphn3 were statistically indistinguishable (Fig. S4C, D), suggesting that the F1041E point mutation selectively disrupts GPCR signaling without altering Lphn3 trafficking or stability.

**Figure 4.**
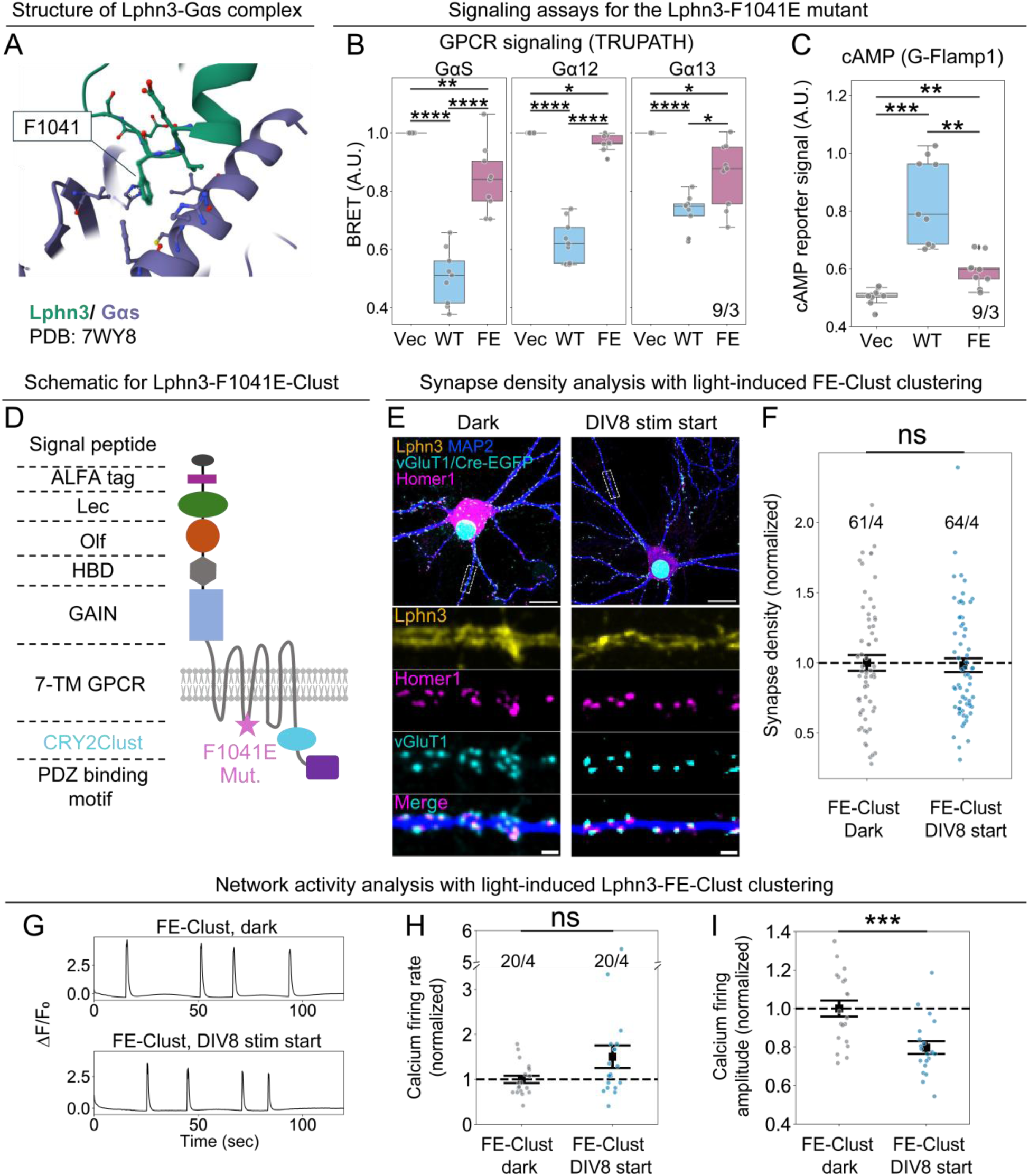
Lphn3 GPCR signaling is essential for clustering-induced synapse formation. **(A)** Structure of the complex of the intracellular 2^nd^ loop of Lphn3 (*green*) with heterotrimeric Gα_S_ protein (*purple*). **(B & C)** G protein coupling assay TRUPATH and cAMP signaling assay in HEK cells expressing vector (Vec), wild type Lphn3 (WT), and Lphn3 bearing the F1041E mutation (FE) with the monomeric mIRFP670^74^ inserted into the C terminal disordered region analogous to Fig. 3A. In **B** and **C**, each gray datapoint represents the BRET readout from an individual well or the average from all cells in a single field-of-view, respectively. (**D**) Domain structure for Lphn3-F1041E-Clust (FE-Clust). **(E)** Representative images of stained neurons expressing FE-Clust. Scale bars: 20 μm (overview)/2 μm (zoom-in). **(F)** Density of excitatory synapses measured by number of puncta positive for both pre- and post-synaptic synaptic markers per μm dendrite. **(G)** Representative population calcium traces. (**H&I**). Synchronous calcium firing rate and amplitude. Mean and SEM are shown in **F, H** and **I**. To account for batch variance, data are normalized to the average value of the dark condition from each batch (denoted as the black dashed line). Dots represent data from each well (**B**), each dendritic segment (**F**) or each field-of-view (**C, H, I**). Statistics: For comparison between more than two conditions, One-way analysis of variance (ANOVA) with post hoc Tukey (equal variance data) or Games-Howell (non-equal variance data) test was used. For comparison between two conditions, independent t-test was used. \*\*\*\**P* < 0.0001; \*\*\**P* < 0.001; \*\**P* < 0.01; \**P* < 0.05; ns: not significant.

We next incorporated the F1041E mutation into the optogenetic clustering construct (Lphn3-F1041E-Clust, Fig. 4D) and tested the role of G protein signaling in clustering-induced synapse formation. Neurons expressing Lphn3-F1041E-Clust were subjected to the same light stimulation protocol as used for WT Lphn3-Clust, beginning at DIV8 and analyzed at DIV14. Upon light-induced Lphn3 clustering, the GPCR signaling-deficient Lphn3 mutant failed to increase excitatory synapse density (Fig. 4E, F). Similarly, network activity as measured by synchronous calcium firing showed no significant increase in firing frequency and a moderate decrease in firing amplitude following clustering (Fig. 4G-I). These results demonstrate that, when uncoupled from downstream GPCR signaling, Lphn3 clustering alone is insufficient to drive synapse assembly but must be coupled to intracellular GPCR signaling.

Given that GPCR signaling is essential for clustering-induced synapse formation, we asked whether clustering directly alters downstream GPCR signaling outputs. Previous work suggests that multimerization of GPCRs, including the adhesion-GPCRs ADGRD1^41^ and ADGRG6^42^, may modulate their signaling activity^43–45^. More broadly, receptor clustering may enhance signaling output by overcoming local inhibitory constraints or by creating a high-intensity local signal that amplifies downstream pathway activation^46–49^. For example, cAMP signaling is compartmentalized by phosphodiesterases (PDEs)^49,50^ and PKA regulatory subunits^51^, limiting the spatial dissipation of the cAMP signal. Since signaling through Gα_S_ is proposed to be the most relevant pathway for Lphn3’s function in synapse assembly^20,21^, we tested whether acute clustering of Lphn3 is sufficient to potentiate GPCR signaling through the GαS–cAMP–PKA pathway.

We co-expressed Lphn3-Clust with the PKA activity reporter Halo-AKAR2.1^52^ in NIH 3T3 fibroblasts (Fig. S5). This reporter operates in the far-red spectrum and is spectrally compatible with CRY2-Clust-based clustering. Upon a brief pulse of blue light to induce Lphn3 clustering, we observed a small increase in PKA activity (Fig. S5B-D). This signal increase was reduced to background levels when the F1041E mutation was introduced into Lphn3-Clust (Fig. S5C&D), indicating that the observed signal depends on intact GPCR coupling. The small global PKA increase suggests that clustering does not elevate the GPCR activity of individual Lphn3 molecules, but rather raises their local density, thereby amplifying cAMP-PKA signaling in confined regions to overcome possible local inhibition^46,49–51^.

In summary, Lphn3 signaling through heterotrimeric G proteins is required for clustering-induced synapse assembly. This may be due to an increase in the local concentration of the high basal signaling activity of Lphn3^20,21,53^ upon clustering, although increased signaling resulting from cluster formation could contribute. A similar mechanism likely applies to related adhesion-GPCRs. Thus, clustering plausibly acts to organize Lphn3 into spatially confined GPCR signaling hotspots that enable effective coupling to downstream effectors.

### Activation of synapse assembly by Lphn3 clustering requires its C-terminal PDZ-binding motif but not its N-terminal FLRT- and teneurin-binding sequences

The function of Lphn3 in organizing excitatory synapse assembly requires its C-terminal PDZ binding motif (PBM)^20^ and extracellular ligand binding domains^19^. The Lphn3 C-terminal PBM binds to postsynaptic density scaffolds such as SHANK^20,54,55^, whereas its extracellular olfactomedin-like (Olf) and lectin-like domains bind to FLRTs and teneurins, respectively^56–59^. Recent *in vitro* reconstitution assays suggested that these interactions function cooperatively: teneurins recruit the presynaptic active zone protein condensate (RIM-BP2/RIM1), whereas Lphn3 recruits the postsynaptic scaffold protein condensate (Shank3/PSD-95/GKAP/Homer3/SynGAP), with heterophilic interactions between Lphn3 and teneurins connecting these pre- and postsynaptic assemblies^20,60,61^. These findings raise the possibility that Lphn3 clustering promotes synapse formation via enhancing Lphn3’s interactions with its presynaptic ligands, recruitment of postsynaptic density scaffolds, or both.

To test these possibilities, we constructed two mutants: a ΔPBM mutant lacking the last three amino acids at the C-terminus that are required for PDZ interactions^20^, and a ΔLecΔOlf mutant which removes both the extracellular teneurin- and FLRT-binding domains of Lphn3^19^ (Fig. 5A). We then tested their ability to support clustering-induced synapse formation.

**Figure 5.**
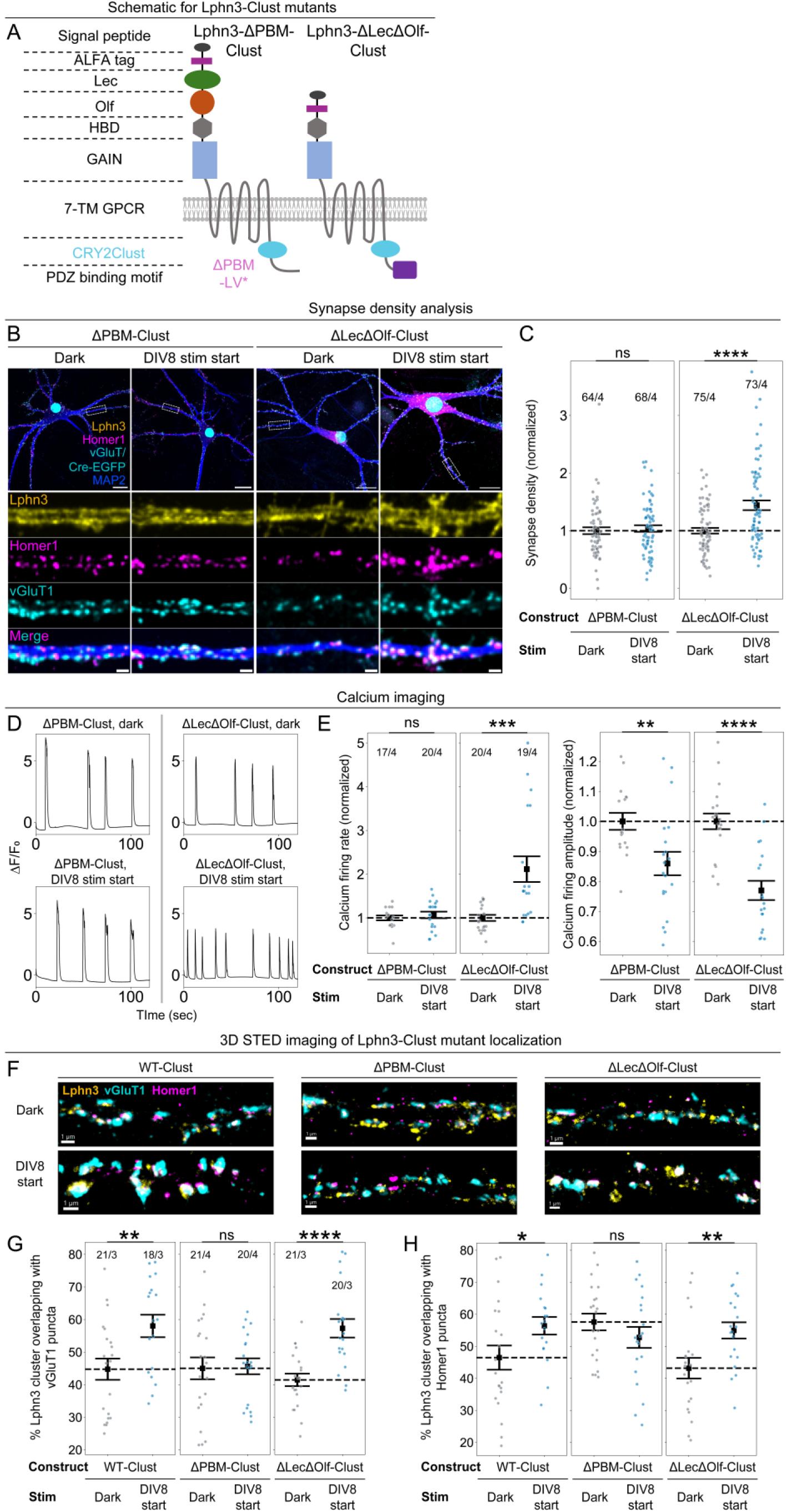
Lphn3 clustering-induced synapse formation requires its PBM but not its teneurin- or FLRT-binding sequences. (**A**) Domain structures of Lphn3-ΔPBM-Clust and Lphn3-ΔLecΔOlf-Clust. **(B)** Representative images of neurons stained at DIV14 for Lphn3, Homer1, and vGluT1. Scale bars: 20 μm (overview) / 2 μm (zoom-in). **(C)** Density of excitatory synapses measured as overlapping vGluT1 and Homer1 puncta. **(D)** Representative calcium imaging traces calculated by averaging the jRGECO1a traces from every cell in a field. **(E)** Quantifications of the synchronous firing rate and calcium signal amplitude. In **C** and **E**, mean and SEM are shown. All data are acquired in paired experimental batches. To account for batch variance, data are normalized to the average value of each batch of a given construct in dark condition, and this average is denoted as a black dashed line. **(F)** Example dendritic segments imaged with 3D STED microscopy displayed in the Imaris 3D view. Scale bars: 1 μm. **(G & H)** Percentage of Lphn3 clusters in cultured neurons overlapping with the synaptic markers vGluT1 (**G**) or Homer1 (**H**). Dots represent data from individual dendritic segments (**C, G, H**) or each field-of-view (**E**). Statistical tests were conducted between dark and light experiments for each construct since these experimental batches are paired. Statistics: independent t-test. \*\*\*\**P* < 0.0001; \*\*\**P* < 0.001; \*\**P* < 0.01; \**P* < 0.05; ns: not significant.

Optogenetic clustering of the ΔPBM mutant failed to induce synapse assembly or enhance synchronous firing in cultured neurons (Fig. 5B-E). A modest reduction in the amplitude of synchronous calcium transients was observed, similar to that obtained with Lphn3-Clust and FE-Clust (Fig. 5E; see Fig. 3J, Fig. 4I).

In contrast to the ΔPBM mutant, optogenetic clustering of the ΔLecΔOlf mutant robustly increased both synapse density and synchronous calcium firing rates (Fig. 5B-E) despite lacking Lphn3’s extracellular ligand-binding domains. This unexpected result suggests that Lphn3 ligand binding is not required for synapse assembly if Lphn3 clustering is induced by light instead of ligand binding. Clustering of both ΔLecΔOlf and ΔPBM mutant Lphn3, like clustering of the signaling mutant construct described above, caused a decrease in the calcium signal amplitude. This decrease may reflect the formation of functionally weaker or less mature synapses, consistent with the inability of these mutants to fully couple receptor clustering to downstream signaling, postsynaptic scaffold assembly, or presynaptic machineries.

To gain further mechanistic insight into how the WT Lphn3 and the ΔLecΔOlf mutant induce synapse formation when artificially clustered, and why the ΔPBM mutant cannot do so, we performed quantitative super-resolution 3D STED imaging. Both ΔPBM-Clust and ΔLecΔOlf-Clust formed membrane-associated nanoclusters (Fig. 5F), indicating that neither interactions through intracellular PBM nor known extracellular ligand-binding domains are individually required for Lphn3 nanocluster formation.

We then investigated whether light-induced Lphn3 clustering alters the ability of Lphn3 mutants to recruit synaptic markers by quantifying the percentage of Lphn3 clusters that overlap with synaptic markers. Driving clustering of both WT and ΔLecΔOlf-mutant Lphn3 increased the fraction of synapse-associated Lphn3 clusters, whereas clustering of ΔPBM-mutant Lphn3 did not (Fig. 5G-H). Notably, the ΔLecΔOlf mutant exhibited an even greater increase in synapse-associated receptors upon induced clustering than WT-Clust (Fig. 5G-H), consistent with a relatively lower baseline (dark) synapse association in the absence of ligand binding. For WT- and ΔLecΔOlf-Lphn3, light-induced clustering did not significantly enhance the sizes of vGluT1 and Homer1 synaptic puncta (Fig. S7D & E), suggesting that Lphn3 clustering enhanced synapse assembly rather than simply enlarging existing synapses. In contrast, light-induced clustering of the ΔPBM mutant resulted in a reduction in vGluT1 and Homer1 puncta size (Fig. S7D & E). These results suggest that the ΔPBM mutant may act in a dominant-negative manner, engaging local GPCR signaling and presynaptic assembly machineries that impose energetic and material demands on the neuron while failing to recruit postsynaptic scaffolds.

To summarize, induction of synapse formation by optogenetic Lphn3 clustering required its interaction with the postsynaptic scaffolds via its PBM, but not its ability to bind to teneurins or FLRTs.

## Discussion

Our findings identify Lphn3 nanoclustering as an essential mechanism that drives Lphn3-induced synapse assembly. Using super-resolution imaging, we show that endogenous Lphn3 assembles into dense nanoclusters containing an average of ∼20 Lphn3 molecules (Fig.1). Lphn3 nanoclusters were observed both in and outside of excitatory synapses with a broad size distribution, but larger nanoclusters containing more Lphn3 molecules were preferentially localized at synapses and the Lphn3 nanocluster size correlated with synaptic strength (Fig. 2).

The correlation between Lphn3 nanocluster size and excitatory synaptic strength suggested that Lphn3 nanoclusters might control synapse assembly. To test this possibility, we employed an optogenetic strategy to acutely induce Lphn3 clustering in neurons. In neuronal cultures, inducing Lphn3 clustering during early network development increased synapse numbers and enhanced network activity, whereas clustering at later stages had little effect. This temporal specificity suggests that Lphn3 clustering is sufficient to drive synaptogenesis but that chronic long-acting Lphn3 clustering is required. Importantly, light-induced clustering of exogenous Lphn3 on a wild-type background in the presence of normal endogenous Lphn3 also caused a dramatic increase (nearly 100%) in synapse numbers (Fig. 3G). Since simply overexpressing Lphn3 does not induce synapse formation beyond normal levels under Lphn3 KO (Fig. 3C, also see^19^), it appears that the extent of Lphn3 nanoclustering instead of the level of Lphn3 is rate-limiting for synapse formation, and that the size of the nanoclusters is critical in inducing synapse formation.

In examining further how Lphn3 clustering promotes synapse assembly, we found that optogenetic clustering of a G protein signaling-deficient Lphn3 mutant failed to enhance synapse formation or increase network activity (Fig. 4). Lphn3 and other synaptic adhesion-GPCRs are characterized by high basal signaling through multiple heterotrimeric G proteins^18,20,39^. Of these, signaling through Gα_S_/cAMP is proposed to be most important for the promotion of synapse formation by Lphn3^20,21^. Clustering of Lphn3 may thus concentrate cAMP signaling to trigger synaptogenesis by inducing the non-linear activation of downstream signaling. Clustering of Lphn3 as a Gα_S_-coupled GPCR may effectively activate synaptogenic signaling through Gα_S_, cAMP, and PKA by overcoming local inhibition imposed by phosphodiesterases and PKA regulatory domains^46,49,50^. Clustering may additionally directly enhance Lphn3 signaling, as has been suggested for other adhesion-GPCRs^41,42^. Consistent with these scenarios, we found that optically induced Lphn3 clustering in NIH 3T3 fibroblasts produced a small increase in total PKA activity (Fig. S5) which could be explained by either an upstream activation of signaling through Gα_S_^20,21,53,62^ or by overcoming an inhibitory threshold. Thus, Lphn3 clustering likely provides a means to generate spatially localized cAMP signal to support synapse formation.

We further dissected the mechanistic requirements for Lphn3 clustering-induced synapse formation in more detail by testing mutants that disrupt intracellular or extracellular interactions. We found that the C-terminal PDZ-binding motif (PBM) of Lphn3, which mediates interactions with postsynaptic scaffolds such as SHANK^54,55^, is essential for clustering-induced synapse formation, consistent with a functional requirement for the PBM in synapse formation^20^. Surprisingly, removal of the Lphn3 lectin-like and olfactomedin-like domains, which mediate the interactions of Lphn3 with presynaptic teneurin and FLRT ligands^19,57^, did not block clustering-induced synaptogenesis. Thus, Lphn3 clustering promotes synapse assembly primarily through intracellular signaling and postsynaptic density scaffold recruitment. Prior studies have shown that the presynaptic Lphn3 ligands teneurin and FLRTs contribute to synapse assembly at least in part by clustering Latrophilins^18,60^. We suggest that optogenetic Lphn3 clustering bypasses the requirement for extracellular ligand binding by effectively performing the same action as these ligands, namely to cluster Lphn3^20,60^. This clustering then causes localized high-intensity cAMP signaling and the massive recruitment of postsynaptic scaffolds to the developing synapse (Fig. 6).

**Figure 6.**
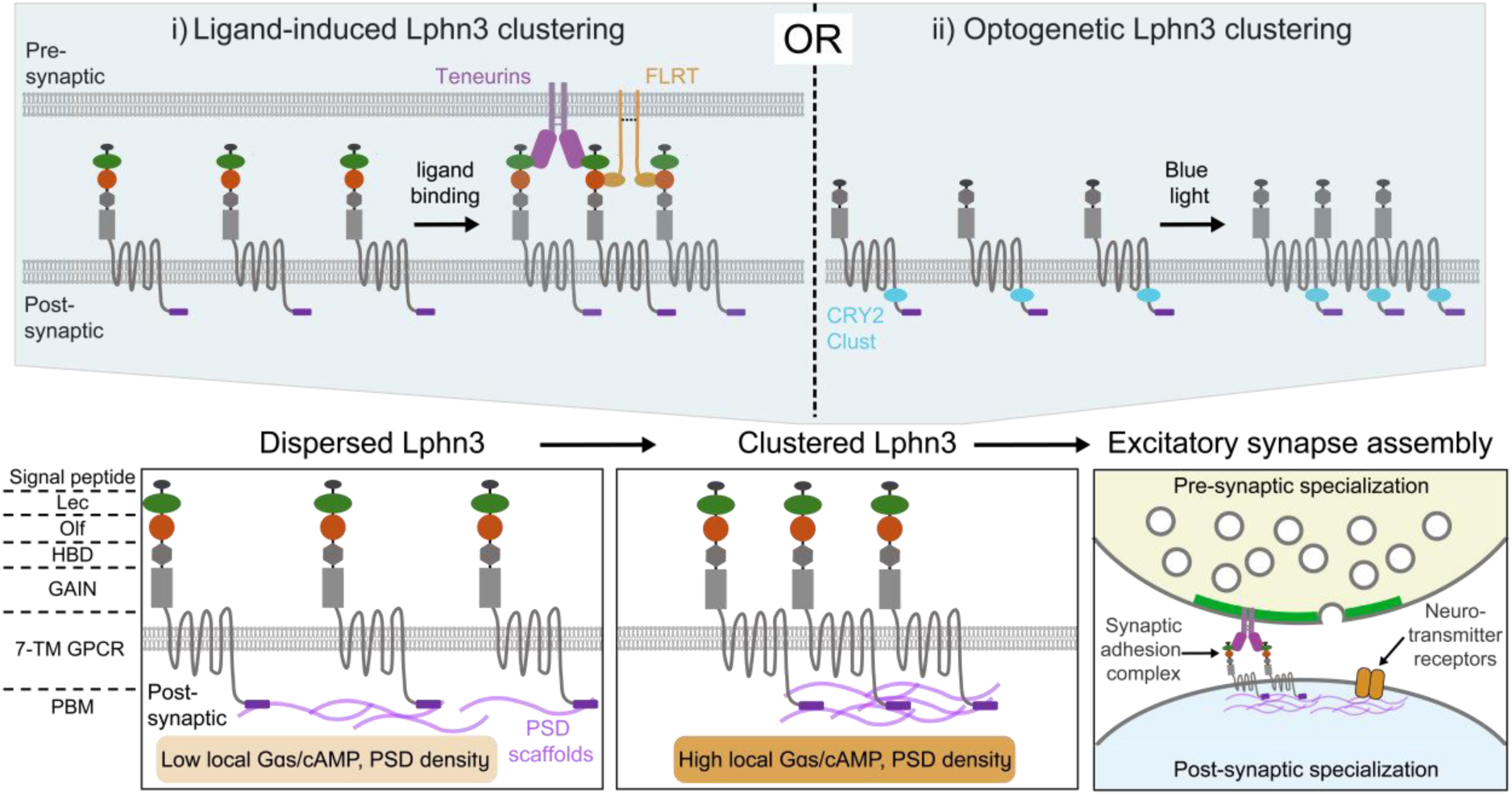
A mechanistic model for Lphn3 clustering during excitatory synapse assembly. Lphn3 clustering can be triggered either by presynaptic ligand binding **(i)** or by optogenetics induction **(ii).** In both cases, clustering increases local Lphn3-associated GPCR signaling and amplifies downstream pathways such as cAMP production. This local signaling enhancement engages post-synaptic scaffold-associated effectors to promote excitatory synapse formation and/or stabilization.

Together, our data thus support a model whereby the spatial organization of Lphn3 serves as a critical regulator of synapse assembly (Fig. 6). In this model, Lphn3 clustering induced by interacting with its presynaptic teneurin and FLRT ligands concentrates local GPCR downstream signaling and recruits PSD-associated effectors to promote the formation and/or stabilization of excitatory synapses. We speculate that the coincident signal from localized cAMP signaling and the recruitment of PSD scaffolds may provide the cues necessary for excitatory synapse formation, an exciting model that could account for the precise co-localization of pre- and postsynaptic specializations at developing synapses^5,6,63^.

Lphn3, like many adhesion-GPCRs, has been proposed to signal through exposure of a tethered agonist following the GAIN domain dissociation^39,53,64,65^. However, a Lphn3 mutant in which GAIN domain autoproteolysis and dissociation is blocked rescued normal synapse formation^19^, indicating that full tethered-agonist exposure is not required for synaptogenesis. Another model of adhesion-GPCR activation is conformational change of the transmembrane region mediated by extracellular ligand binding and/or mechanical force^66,67^. Our findings support an additional, alternative general mechanism of adhesion-GPCR activation whereby extracellular ligand binding drives receptor clustering, thereby increasing local GPCR-associated signaling (Fig. 6) and enabling neurons to control signaling output through the spatial organization of Lphn3 rather than solely through conformational change of the receptor. Given the widespread expression of adhesion-GPCRs in the central nervous system, as well as their involvement in multifarious neurodevelopmental and neurological disorders^15^, similar organizational principles may operate for other adhesion-GPCRs in other physiological and disease contexts.

Beyond these biological insights, the optogenetic clustering strategy developed here provides a generalizable approach to manipulate synaptic receptor organization, and hence synapse formation, with spatial and temporal precision. Previous optogenetic approaches have successfully controlled synaptic function by manipulating postsynaptic density components or neurotransmitter receptors^68,69^. Our work extends these concepts to directly regulate the nanoscale assembly of a synaptic adhesion-GPCR and shows that this provides a viable means to control synaptogenesis. Enabling direct control over receptor assembly provides a potentially powerful way to dissect how receptor organization shapes signaling and function in synapses and other cellular systems.

This study has several limitations. First, we focused on an abundant splice variant of a single Latrophilin isoform with established roles in excitatory synapse formation^19,20^. Whether similar mechanisms operate across other Latrophilins or other splice variants of Lphn3 remains unknown. Second, although our results support a role for Lphn3 clustering in synapse formation, our experiments cannot tell whether Lphn3 clustering initiates formation of nascent synapses or stabilizes immature or mature synapses. The fact that light-induced clustering from DIV12-14 has little effect on synapse numbers (Fig. 3) supports a role in synapse initiation, but rapid normal turnover of at least some synapses^70,71^ makes it impossible to distinguish between synapse initiation and maintenance. Third, while this work centers on synapse formation, Latrophilins contribute to other developmental processes^1,72,73^ and the role of the spatial organization of Lphn3 in these other contexts remains unexplored. Fourth, although clustering moderately activates signaling downstream of Gα_S_ in heterologous cells, we have not directly tested whether it similarly elevates Gα_S_-mediated signaling in neurons, where the signaling dynamics and regulatory constraints may differ. This experiment is technically challenging because it requires high expression levels of signaling reporters, which are difficult to achieve in neurons and may compromise neuron health. Fifth, all of our neuronal experiments were performed in primary cultures and it is possible that the in vivo functions of Lphn3 are mechanistically different, although this seem unlikely given that in all previous studies, the in vivo and culture results for Lphn3 perfectly aligned^19–21^. Finally, although GPCR signaling and the PDZ-binding motif are required for clustering-induced synapse formation, multiple downstream effector proteins linking Lphn3 clustering to synapse assembly remain unidentified and represent an important direction for future work.

Despite these limitations, this study identifies nanoscale clustering as a central mechanism by which Lphn3 as a classical adhesion-GPCR induces synapse assembly. Given the broad expression patterns and strong links to neurodevelopmental disorders of adhesion-GPCRs^15,16^, these insights provide a framework for understanding disease mechanisms involving adhesion-GPCRs and for developing strategies to therapeutically modulate adhesion-GPCR function.

## Acknowledgements

We would like to thank Stanford Cell Sciences Imaging Facility (CSIF) and Stanford OHNS imaging facility for imaging support, particularly David Lenzi and Katharine Miller for their technical assistance with microscopy. We are also grateful to colleagues who provided valuable discussions, in particular Silas Boye Nissen, Daniel Matus, Shaoyuan Zhu, Xuzhong Yang, Xuchen Zhang, Shuai Wang, the members of the Dunn lab, and the members of the adhesion molecules subgroup in the Südhof lab.

## Declaration of generative AI and AI-assisted technologies in the writing process

During the preparation of this work, the authors used ChatGPT and Gemini in order to improve the readability and language of the manuscript. After using these tools/services, the authors reviewed and edited the content as needed and take full responsibility for the content of the published article.

## Author contributions

C.H. performed all data acquisition and analysis except for the TRUPATH assay which was performed by A.B.; A.B. also performed data acquisition with cultured neurons. G.L. with C.H. performed single molecule counting experiments and data analysis; A.R.D. and T.C.S. acquired funding for this project; C.H., A.R.D., and T.C.S. conceptualized the project, designed the experiments, wrote and edited the manuscript with contribution from all authors.

## Materials & Correspondence

Correspondence and requests for materials and/or software should be addressed to Alexander R. Dunn.

## Data availability

All raw data and analysis code used in this study will be deposited to the Stanford Data Repository by the time of publication.

