## Supplementary Figures for "Clustering of the Adhesion-GPCR Latrophilin-3 Drives Excitatory Synapse Assembly"

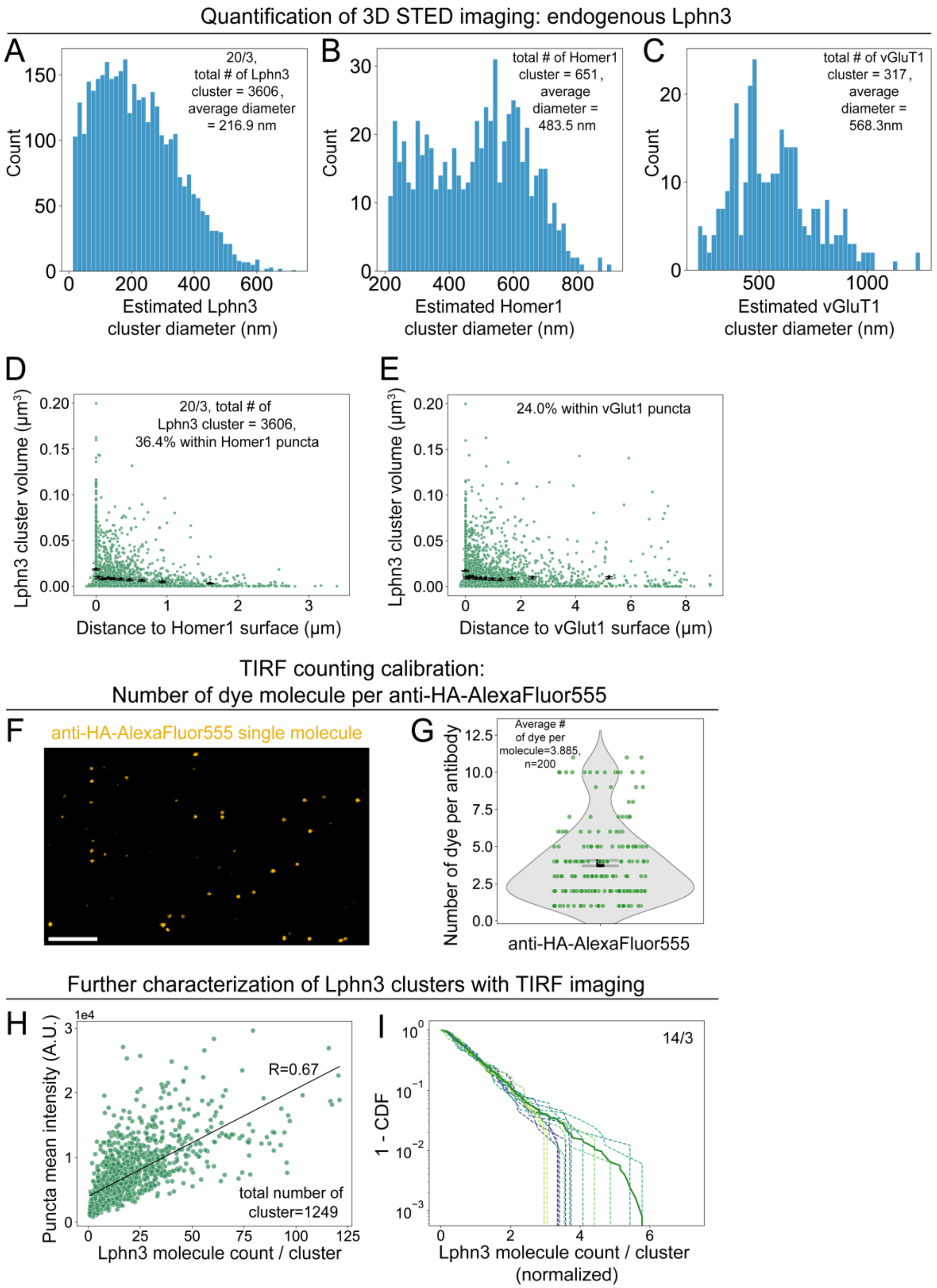


**Figure S1**. Further characterization of 3D STED and TIRF imaging of endogenous Lphn3**. (A-C)** Distribution of cluster diameters quantified from 3D STED for endogenous Lphn3 (**A**), Homer1 (**B**), and vGluT1 (**C**). Diameters are estimated from cluster volumes assuming perfect spheres. **(D&E)**

Scatter plots of the Lphn3 cluster volume vs. distance to postsynaptic Homer1 (**D**) and presynaptic vGluT1 (**E**) surfaces. Data points less than or equal to zero denote Lphn3 clusters located within the volume defined by synaptic markers. Data are organized into 14 equal-size bins. Within each bin, black dots show average, while error bars show the 95% confidence interval from bootstrap resampling (1000 iterations). (**F**) Representative single-molecule TIRF images of the anti-HA antibody (conjugated with AlexaFluor555) used to stain neurons. Scale bar: 2 μm. (**G**) Distribution of AlexaFluor555 dye molecules per antibody. Error bar shows mean ± SEM. (**H**) Number of Lphn3 molecule per cluster calculated from TIRF bleaching experiments plotted against Lphn3 cluster mean intensity. A linear fit and correlation coefficient are displayed. (**I**) Complementary cumulative distribution function (1-CDF) of Lphn3 molecule count per cluster, normalized to the means of each dendritic segment. 1 – CDF of each dendritic segment is shown as colored dashed lines, and the average of all dendritic segments is shown as the solid green line. The 1-CDF curves resemble exponential distributions.

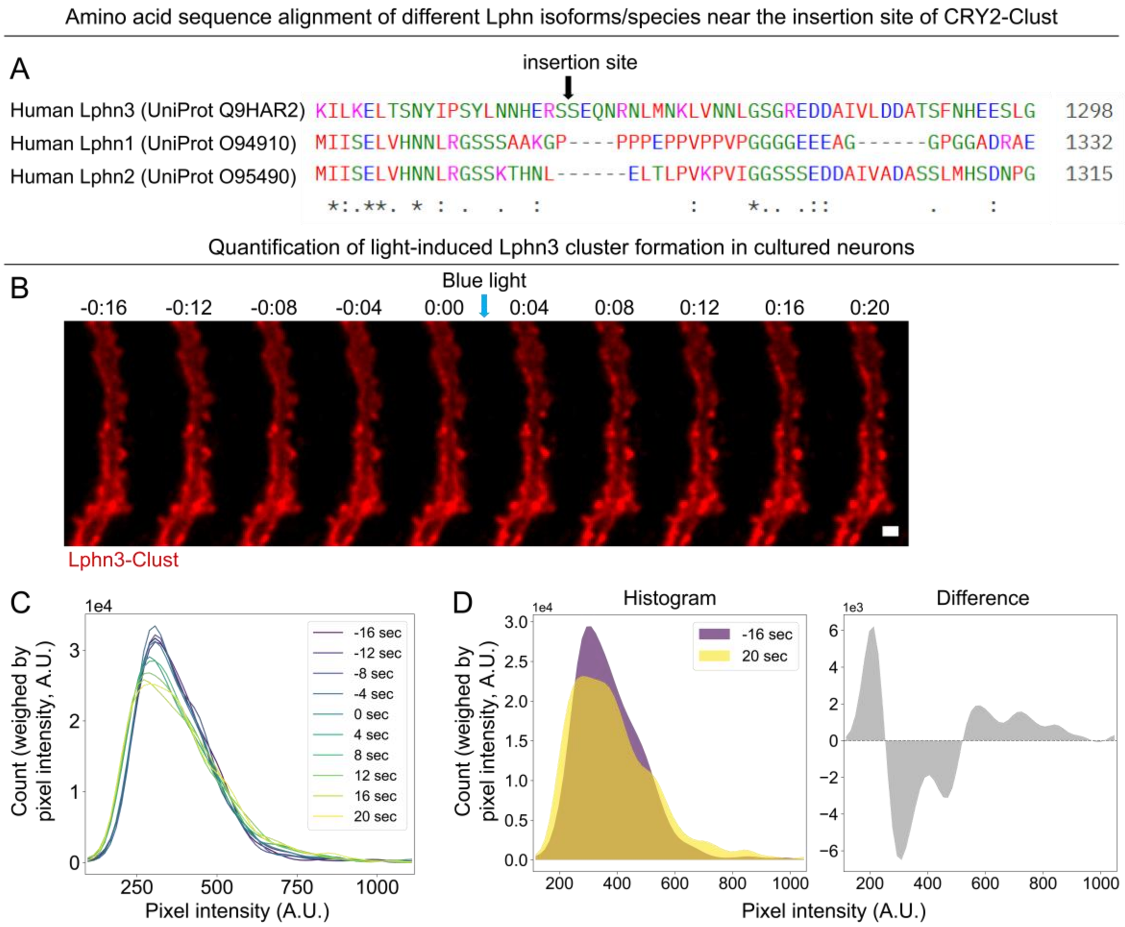


**Figure S2. Insertion site of CRY2Clust in the C-terminal cytoplasmic Lphn3 sequences and further quantifications of Lphn3 opto-clustering in cultured hippocampal neurons (A).** Multiple sequence alignment of human Lphn1, Lphn2, and Lphn3 around the insertion site of CRY2-Clust in the C terminal disordered region. Alignment conducted using Clustal Omega[38] (asterisk = fully conserved residue; colon = strongly conserved residue; period = weakly conserved residue). Fluorescent protein (mVenus) insertion in the analogous site retained Lphn3's ability to activate cell signaling pathways associated with actin-cytoskeleton remodeling[39] as well as its ability to couple to G proteins[20] (**Fig. 4B**). Throughout this work, rescue constructs were constructed from human Lphn3. **(B).** Time series of Lphn3-Clust in live cultured neurons, the same series as in **Fig. 3B**. Lphn3-Clust was surface labeled with AlexaFluor647-conjugated ALFA nanobody. Blue light stimulation was applied immediately after 0:00 sec. Scale bar: 5 µm. **(C & D).** Weighed histograms of Lphn3 pixel intensity for the time series (weighed by pixel intensity). Histograms were extracted from masked dendritic areas. **C** shows histograms for each time point, whereas **D** shows the histograms before (*purple*) and after (*yellow*) light stimulation (*left*) and the difference between the two histograms (*right*). Blue light led to an increase in high-intensity values (intensity ~700), an increase in low intensity values (intensity ~200), and a decrease of moderate intensity values (intensity ~300). These changes correspond to the enhancement of signal at pre-existing clusters, the increased area with low signal, and the reduction of diffuse Lphn3 signal, respectively.

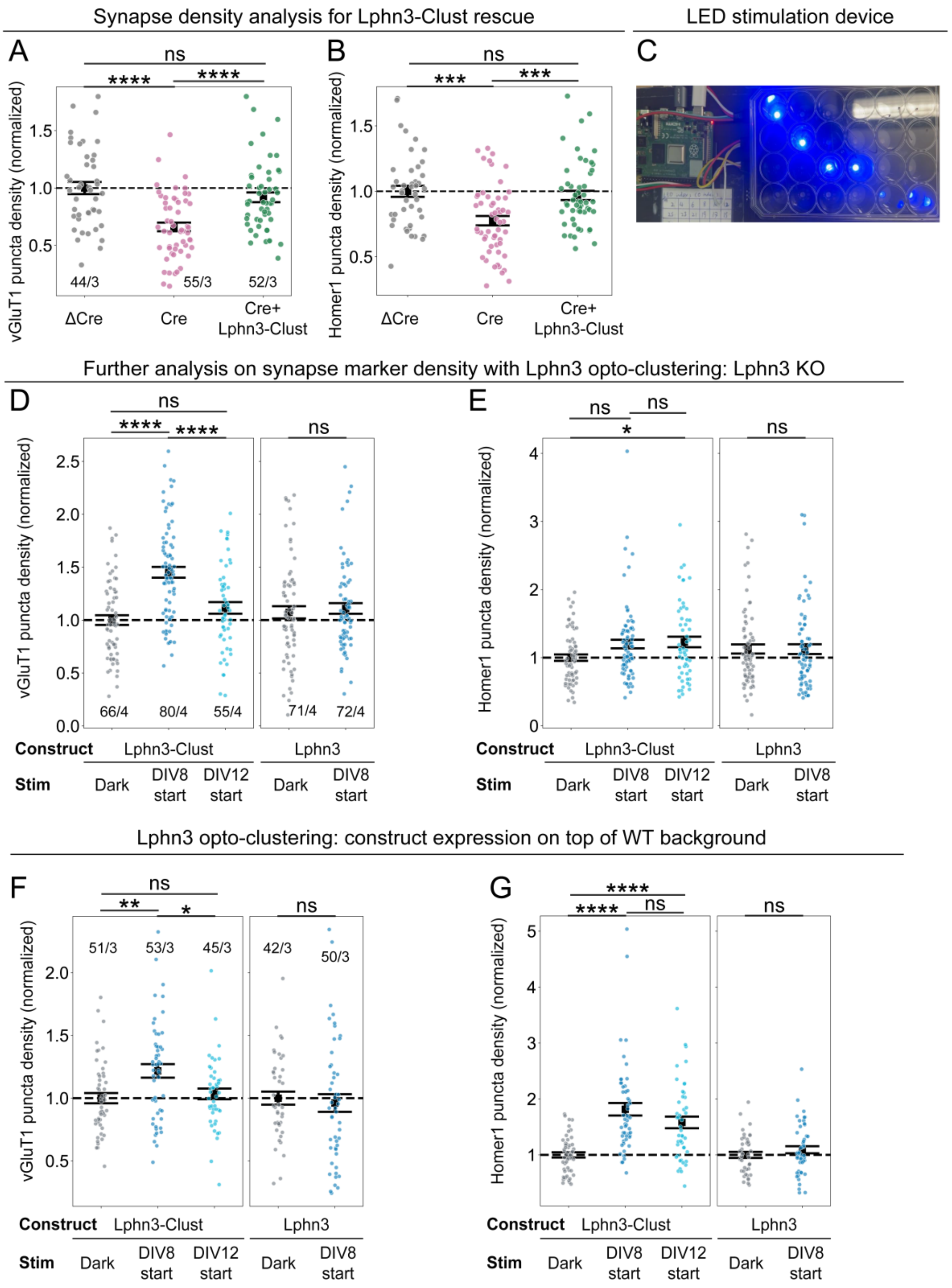


**Figure S3. Further measurements of Lphn3 opto-clustering in cultured hippocampal neurons (A & B).** Measurement of vGluT1 and Homer1 marker puncta density with Cre (Lphn3 knockout), ΔCre (knockout control), and Cre followed by rescue with Lphn3-Clust. To account for batch variance, data are normalized to the average value of each batch of the ΔCre condition, and this average is denoted as black dashed line at one. The same dataset as **Fig. 3C**. **(C)** Photo of the LED blue light stimulation device. **(D & E).** Measurement of presynaptic marker (vGluT1) and post-synaptic marker (Homer1) density in DIV14. The same dataset as in **Fig. 3F.** To account for batch variance, data are normalized to the average value of each batch of the Lphn3-Clust dark condition, and this average is denoted as black dashed line at one. **(F & G).** Measurement of vGluT1 marker puncta density, and Homer1 marker puncta density, with construct expression in wild-type neurons from CD1 pups. To account for batch variance, data are normalized to the average value of each batch of dark conditions, and this average is denoted black dashed line at one. Statistics: For comparison between more than two conditions, One-way analysis of variance (ANOVA) with post hoc Tukey (equal variance data) or Games-Howell (non-equal variance data) test was used. For comparison between two conditions, independent t-test was used. ****$P$ < 0.0001; ***$P$ < 0.001; **$P$ < 0.01; *$P$ < 0.05; ns: not significant.

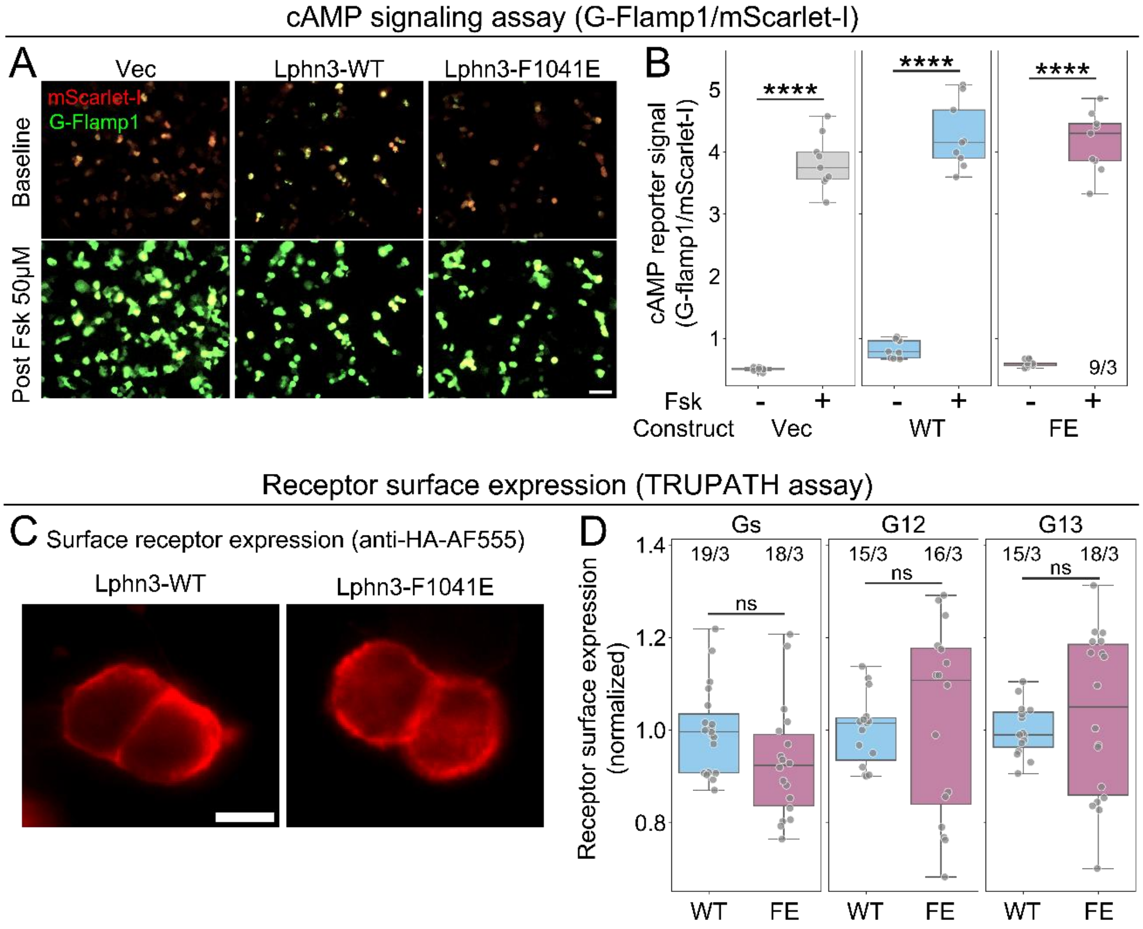


**Figure S4**. **Further characterization of cAMP and TRUPATH signaling assays in heterologous cells.** (**A**) cAMP signaling assay in HEK cells. HEK cells co-expressed the cAMP reporter mScarlet-I-G-Flamp1 and vector (Vec)/ WT-Lphn3 (WT)/ Lphn3-F1041E (FE). Shown are representative images right before and 15 minutes after addition of 50 µM forskolin (Fsk). Scale bar: 100 µm. (**B**) Quantification of the cAMP signal before and after forskolin addition for cells expressing different receptors, demonstrating the G-Flamp1 reporter functions properly. Grey datapoints and box plots show the averages from individual field-of-views. (**C**) Representative widefield images of surface-labeled Lphn3-WT and Lphn3-F1041E expressed in HEK cells. Cells were live-labeled with anti-HA-AlexaFluor555 before fixation. Scale bar: 10 µm. (**D**) Quantification of surface receptor expression from AlexaFluor555 intensity in the same batch of HEK cells from the TRUPATH assay. For each transfection, expression levels were normalized to the average of the corresponding Lphn3-WT levels. Grey dots and box plots show the averages from individual field-of-views. All images in the same panel are shown with the same brightness and contrast settings. Receptor constructs all have an N-terminal HA tag and the monomeric mIRFP670 fluorescent protein inserted in the C terminal disordered region (analogous to insertion site of CRY2Clust in **Fig. 3**). Statistics: independent t-test between field-of-views was used. Ns: not significant, ****$P < 0.0001$.

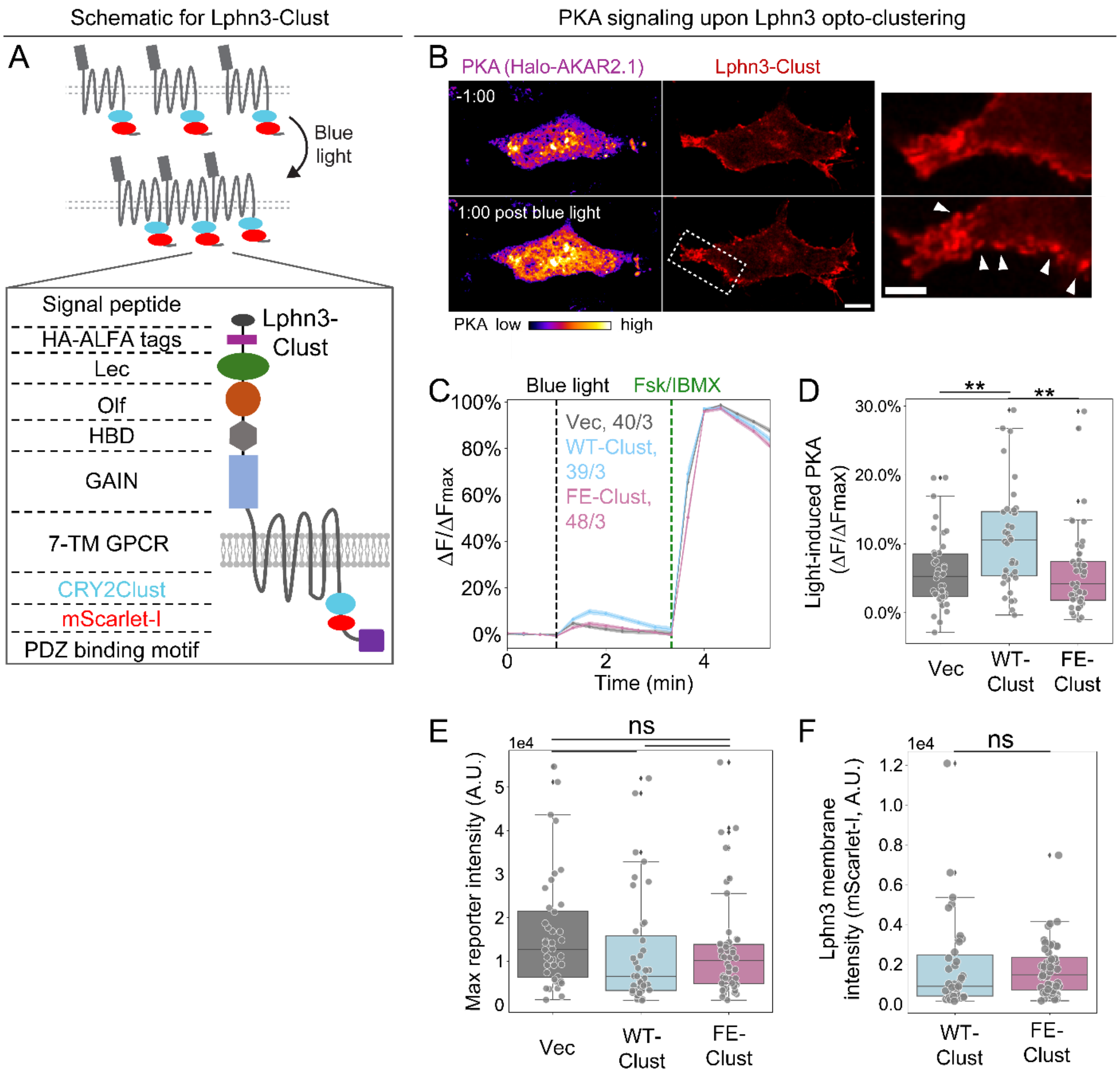


**Figure S5. Constitutive GPCR signaling by Lphn3 involving downstream G$\alpha_S$ signaling is slightly elevated by optogenetic Lphn3 clustering.** (**A**). Schematic of the light-inducible Lphn3 clustering strategy and domain structure of Lphn3-Clust. An optogenetic domain CRY2Clust and monomeric fluorescent protein mScarlet-I are inserted into the same cytosolic region as in **Fig. 3**. (**B**). Images of an NIH3T3 fibroblast cell co-expressing the PKA reporter Halo-AKAR2.1 Lphn3-Clust, one minute before and after 488 nm light stimulation. *Right*: enlarged images of the Lphn3 channel. White arrow heads point to regions where surface Lphn3 form clusters upon light stimulation. Scale bars: 10 μm (cell overview), 5 μm (zoom-in). NIH3T3 fibroblast cells were chosen for this assay for their flat morphology and low transfection efficiency, both of which facilitate Lphn3 cluster visualization and low levels of basal receptor clustering. (**C**). Temporal traces of the normalized PKA reporter activity upon blue light stimulation in NIH3T3 fibroblast cells. Cells were co-transfected with vector (Vec), wild-type Lphn3-Clust (WT-Clust), and Lphn3-Clust bearing the F1041E mutation (FE-Clust), Traces show mean ± SEM calculated from individual cells. (**D**)**.** Quantification of light-induced PKA activity increase (maximum increase from baseline) upon blue light stimulation for the three constructs, extracted from **C**. Each grey datapoint represents a cell, while the box plot shows data distribution. **(E)** Maximum PKA reporter intensities induced by forskolin and IBMX are comparable for all three constructs. (**F**). Lphn3 membrane expression in the PKA signaling assays, measured by mScarlet-I average intensity extracted from membrane masks. All images in the same panel are shown with the same brightness and contrast settings. Statistics: For comparison of more than two conditions, one-way analysis of variance (ANOVA) with post hoc Tukey (equal variance data) or Games-Howell (non-equal variance data) tests was used. For comparison of two conditions, independent t-test was used. ****$P$ < 0.0001; ***$P$ < 0.001; **$P$ < 0.01; *$P$ < 0.05; ns: not significant.

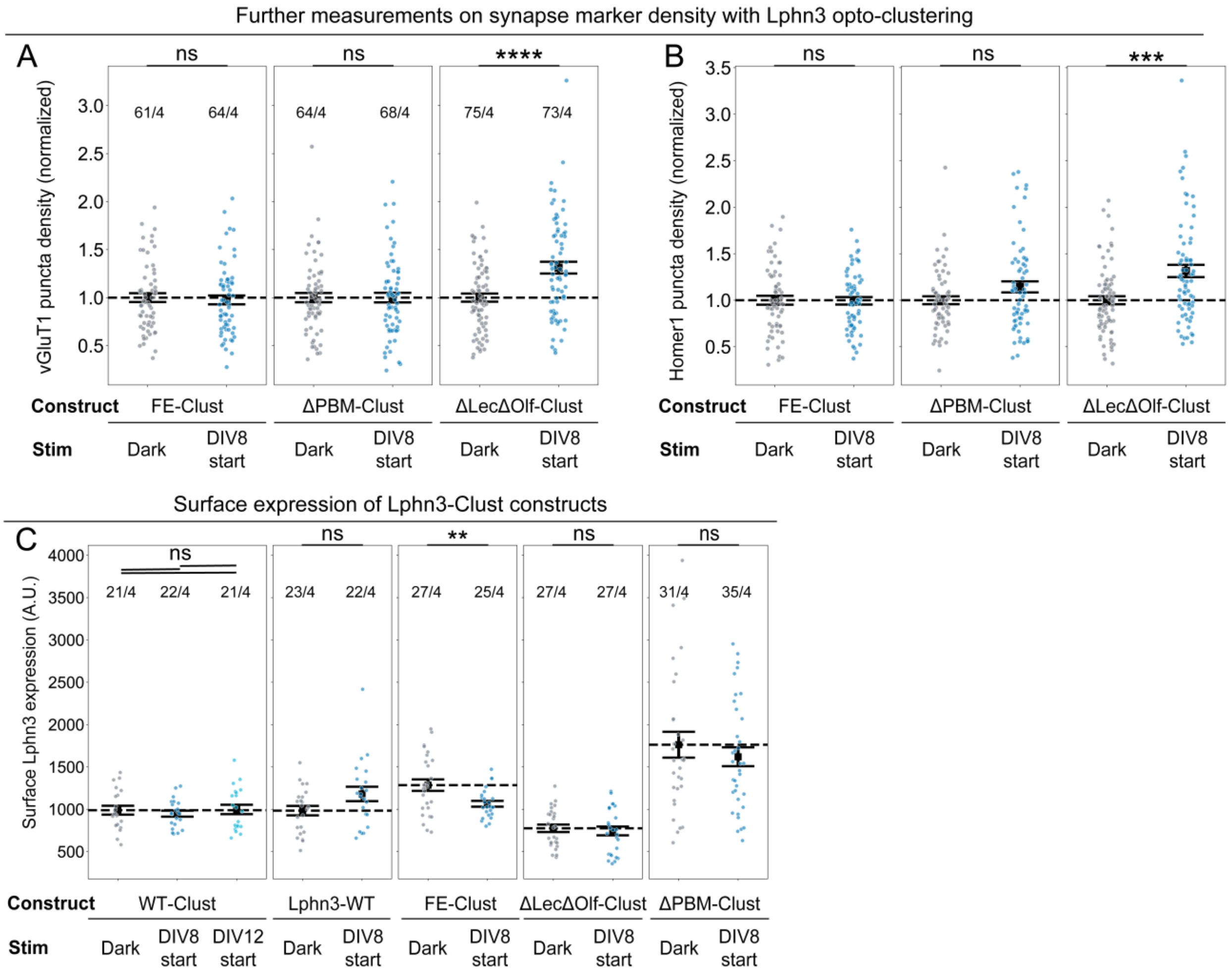


**Figure S6**. **Further characterization for Lphn3-Clust mutants in neurons. (A and B).** Measurement of presynaptic marker (vGluT1) and post-synaptic marker (Homer1) density for fixed and stained DIV14 neurons. The same dataset as in **Fig. 4F and Fig. 5C.** To account for batch variance, data are normalized to the average value of each batch from the dark condition, and this average is denoted as black dashed line at one. (**C**) Surface expression of Lphn3-Clust constructs in cultured neurons under different conditions. Dots represent data from each dendritic segment (**A & B**) or each field-of-view (**C**). Statistics: independent t-test. ****$P$ < 0.0001; ***$P$ < 0.001; **$P$ < 0.01; *$P$ < 0.05; ns: not significant.

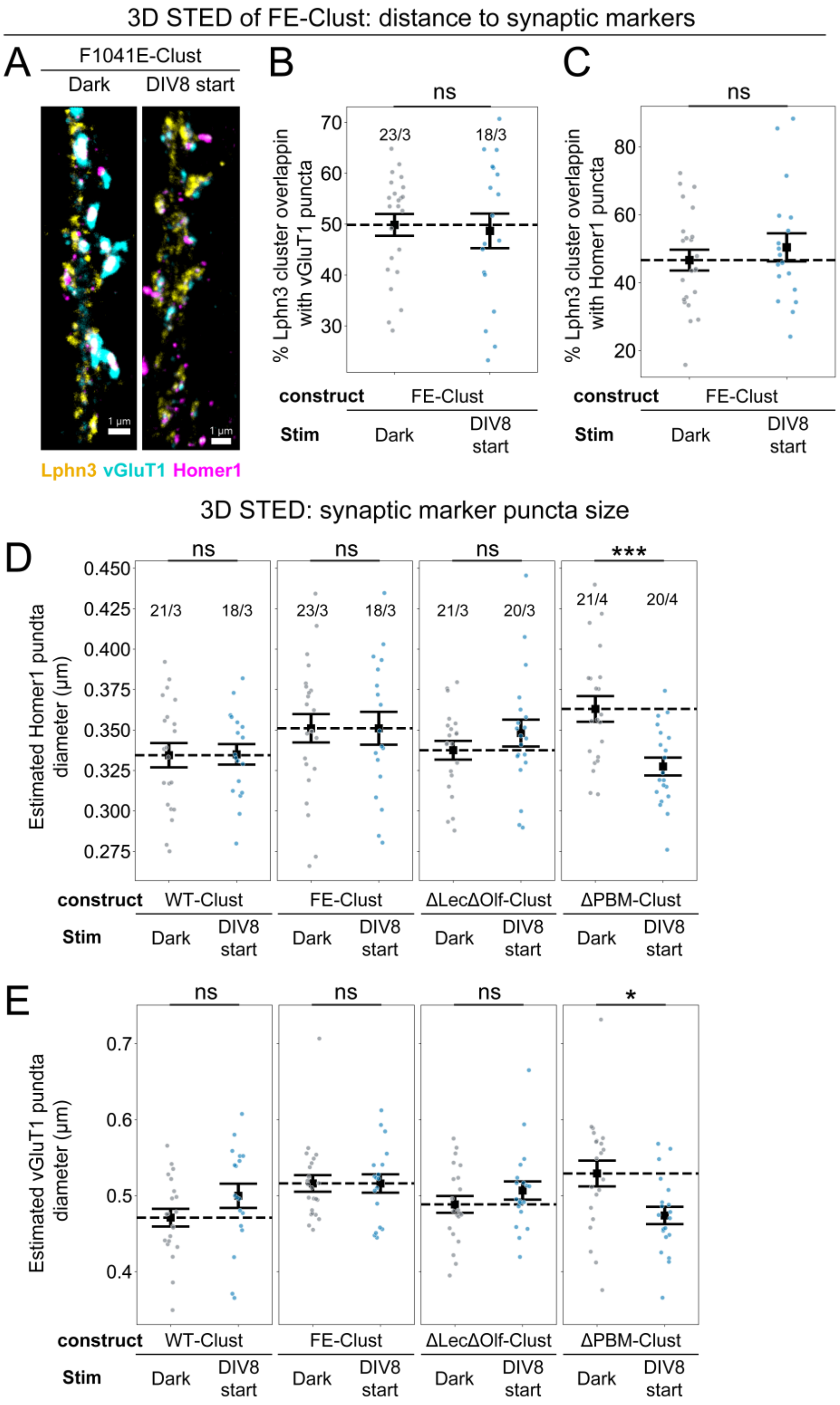


**Figure S7**. **Super-resolution 3D STED images of Lphn3-Clust in cultured hippocampal neurons. (A).** Example dendritic segment expressing Lphn3-F1041E-Clust construct imaged with 3D STED microscopy shown in Imaris volume view. Scale bars: 1 µm. **(B&C).** Percentage of Lphn3-FE-Clust clusters that overlap with vGluT1 puncta (**B**) or Homer1 puncta (**C**). **(D-E).** Estimated puncta diameter of Homer1 (**D**) and vGluT1 (**E**). Puncta diameters were calculated by treating each puncta as a perfect sphere. Note that the Homer1 (**D**) and vGluT1 puncta diameters (**E**) observed with the Lphn3-ΔPBM mutant in darkness appear to be larger than those of the other

Lphn3 mutants and that this apparent increase is reversed by light-induced clustering, could suggest that the absence of PSD95 and/or SHANK binding by Lphn3 causes an increase in synapse volume that is reversed by Lphn3 clustering. Note though that this conclusion cannot be firmly made since the diameter measurements for the different mutants were performed at different times and only dark vs. light conditions were performed in parallel and can be directly compared. Statistical tests were conducted between dark and light experiments for each construct since those experiments are paired. Dots represent data from each dendritic segment. Statistics: independent t-test was used. ***$P < 0.001$; *$P < 0.05$; ns: not significant.

**Supplemental videos (legend)**

**Video S1**. Blue light stimulation induces rapid Lphn3-Clust clustering in cultured neurons. Scale bar: 5 µm. A 2-second pulse of blue light was applied after 0 second.

**Video S2**. A device to apply different light stimulation conditions to different wells of a 24-well plate.
