## Supplementary Methods for "Clustering of the Adhesion-GPCR Latrophilin-3 Drives Excitatory Synapse Assembly"

#### Mouse handling

Mutant Lphn3 conditional knock-in mice containing N-terminal HA tag were generated previously[1] and used for all neuron experiments. Mice were kept at the Stanford Veterinary Service Center, on a 12-hour light-dark cycle and food and water provided ad libitum. Newborn mice were weaned at postnatal days (P) 18-21 and housed in groups with less than 5 mice per cage. All animal procedures conform to the National Institutes of Health Guidelines for the Care and Use of Laboratory Animals and were approved by the Stanford University Administrative Panel on Laboratory Animal Care.

Primers for genotyping are:

Forward: ATTGCTTATGTATCAGTTTGGC

Reverse: GTATACACCACTAGGAAGACTG

#### Plasmids and viruses

| **Plasmid** | **source** |
|---|---|
| TRUPATH assay plasmids | Addgene, #1000000163 |
| pCAG-G-Flamp1 | Addgene, #188567 |
| ePBCAG-bsd-mScarlet-I-G-Flamp1 | Custom made with Gibson cloning. |
| pcDNA3.1-HaloAKAR2.1-T2A-EGFP | Addgene, #216489 |
| pGL4.29-CRE-minP-*luc2P* (CRE transcription reporter) | Gift from Dr. Stephan C. Blacklow at Harvard |
| pRL-TK (*Renilla* luciferase) | Promega |
| ePBCAG-IgK-HA-ALFA-hsLphn3-CRY2Clust-mScarlet(i)-i_SS1+SS2_+SS5_+SS7-hUBC-Puro | Custom made with Gibson cloning. |
| ePBCAG-IgK-HA-ALFA-hsLphn3-F1041E-CRY2Clust-mScarlet(i)-i_SS1+SS2_+SS5_+SS7-hUBC-Puro | Custom made with Gibson cloning. |
| ePBCAG-hUBC-Puro-MCS-Expand | Gift from Hiromitsu Nakauchi's lab at Stanford. |
| ePBCAG-IgK-HA-ALFA-hsLphn3-mIRFP670 | Custom made with Gibson cloning. |
| ePBCAG-IgK-HA-ALFA-hsLphn3-F1041E-mIRFP670 | Custom made with Gibson cloning. |

| **Viral plasmid** | **source** |
|---|---|
| Lenti-hSyn-NLS-EGFP-ΔCre | Custom made with Gibson cloning. |
| Lenti-hSyn-NLS-EGFP-Cre | Custom made with Gibson cloning. |
| FSW-iGluSnFR3.857.GPI | Addgene, #205986 |
| Lenti-hSyn-FSW-jRGECO1a | Gift from Dr. Shuai Wang at Stanford. |
| Lenti-hSyn-IgK-ALFA-hsLphn3 | Custom made with Gibson cloning. |
| Lenti-hSyn-IgK-ALFA-hsLphn3-CRY2Clust | Custom made with Gibson cloning. |

| | |
|---|---|
| Lenti-hSyn-IgK-ALFA-hsLphn3-F1041E-CRY2Clust | Custom made with Gibson cloning. |
| Lenti-hSyn-IgK-ALFA-hsLphn3-ΔLecΔOlf-CRY2Clust | Custom made with Gibson cloning. |
| Lenti-hSyn-IgK-ALFA-hsLphn3-CRY2Clust-ΔPBM | Custom made with Gibson cloning. |

### Statistics

The numbers of technical and biological replicates are indicated for each dataset in the corresponding figure. Mean and SEM from technical replicates are shown in plots unless specified otherwise. Statistical tests were conducted with technical replicates unless specified otherwise, using the Pingouin, SciPy.stats, and Statannot[2] Python packages. For comparison between two conditions, a two-tailed Student's t-test assuming equal variances was used unless otherwise specified. For comparison between more than two conditions, one-way analysis of variance (ANOVA) with post hoc Tukey (for equal variance data) or Games-Howell (for unequal variance data) test was used. ****$P < 0.0001$; ***$P < 0.001$; **$P < 0.01$; *$P < 0.05$, ns: not significant.

### Primary hippocampal neuron culture

Primary neurons from the hippocampi of neonatal mice were prepared as previously described[1,3]. In short, DIV0-1 Lphn3 conditional mouse pups or CD1 mouse pups were dissected in cold HBS to extract hippocampi. Tissues were digested in 1% (v/v) papain (Worthington) for 16 minutes at 37 °C, washed twice in warm HBS, then once in plating medium (MEM containing 10% FBS, Sigma F0926, and 2 mM GlutaMAX), followed by gentle pipetting in plating medium to break apart the cells. Cells were filtered through a 70-μm cell strainer before seeded onto #1.5 glass surfaces pre-coated by Matrigel (Corning). Cells harvested from 5 pups were seeded into two 24-well plates with black wall (Cellvis P24-1.5H-N). 14-18 hours post seeding, 95% of the plating media was replaced from each plate with neuron maintenance medium (Neurobasal A with 5% FBS, Sigma F0926, 2% Gem21 NeuroPlex Supplement, and 2 mM GlutaMAX). Cells were maintained in a humidified incubator at 37 °C under 5% $CO_2$. At DIV 3, 7, and 11, 50% of the media was replaced, during which 2μM Ara-C was supplemented. When indicated, Cre/ΔCre viruses were added at DIV3, rescue Lphn3 viruses were added at DIV4, and jRGECO1a/ iGluSnFR3 viruses were added at DIV7. For cells transduced with optogenetic constructs, plates were kept in the dark before harvested for assays at DIV14.

### Culturing HEK293T and 3T3 cells

HEK293T cells were cultured in DMEM growth medium (Gibco 12430054) supplemented with 1% penicillin-streptomycin (Gibco 15140148), 10% fetal bovine serum (Corning 35-011-CV) and 1% sodium pyruvate (Gibco 11360070). Mouse NIH-3T3 fibroblasts were cultured in DMEM growth medium (Gibco 12430054) supplemented with 1% penicillin-streptomycin (Gibco 15140148), 10% bovine calf serum (Sigma 12138C) and 1% sodium pyruvate (Gibco 11360070). The medium was renewed 2-3 times a week, and cells were passaged before they reached 80% confluency. Cells were maintained in a humidified incubator at 37 °C under 5% $CO_2$

### Opto-stimulation device construction and opto-stimulation

The opto-stimulation device was built using WS2812B RGB LED strips (BTF-Lighting) controlled by a Raspberry-Pi. Individual LEDs were programmed using the Adafruit CircuitPython NeoPixel library. Adapters for 24-well plates were custom-designed in Shapr3D and fabricated with black ABS plastic on a Creality Ender-3 3D printer. Opto-stimulation applied to primary neuron culture consisted of 2 second pulses of blue LED light at 100% brightness every 5 minutes.

### Lentivirus preparation

Lentivirus stocks were generated by transfecting HEK293T cells (ATCC) with the second-generation packaging system. HEK293 cells were seeded in T25 flasks, grown to 50% to 60% confluency, and transfected using the calcium phosphate transfection method with the following plasmids: 3.6μg of the shuttle vector together with helper plasmids (1μg pRSV-REV, 2.5μg pMDLg/pRRE, and 1.3μg vesicular stomatitis virus G-protein). DNA was diluted in 250 uL $ddH_2O$ with 0.25 mM $CaCl_2$ before being added dropwise to 250 uL of 2× HEPES-buffered saline (274 mM NaCl, 10 mM KCl, 1.5 mM Na2HPO4, 12 mM glucose, 42 mM HEPES, pH adjusted to 7.05 with NaOH) followed by brief vortexing. After a 15-minute incubation at room temperature, the mixture was added dropwise to HEK293T cells. 24 hours after transfection, medium was replaced by 5mL neural maintenance medium. After an additional 24 hours, virus-containing medium was harvested, cell debris was removed by centrifugation at 2000 x g for 3 minutes, and the virus-containing supernatant was filtered through a 0.22μm filter before being stored in aliquots at -80°C. Virus titers were determined as follows. For the Cre/ΔCre virus, the lowest amount of virus that achieved 100% transduction efficiency (through nuclear GFP signal) was used. Efficient Cre-mediated knockout was confirmed by surface labeling of endogenous Lphn3 using a rabbit anti-HA antibody (Alexa Fluor® 555 Conjugate, Cell Signaling # 55420, detailed methods see *Immunohistochemistry* below).

To equalize the expression levels of Lphn3 rescue constructs in neurons, virus titers from different mutants were measured by quantifying surface expression levels of ALFA-Lphn3 constructs through surface staining using the FluoTag-X2 anti-ALFA nanobody (Alexa647 conjugated, NanoTag, details, see *Immunohistochemistry* for details).

### Immunohistochemistry

Surface labeling was conducted on live cells (primary neurons, HEK cells, and NIH-3T3 fibroblasts) in culture medium inside a 37 °C incubator with 5% $CO_2$ for 30-45 minutes before fixation with 4% PFA in PBS for 10 minutes at 37 °C. Antibodies used for surface labeling are: FluoTag-X2 anti-ALFA nanobody (Alexa647 conjugated, NanoTag, 1:400), FluoTag®-X2 anti-ALFA nanobody (Abberior STAR 635P conjugated, NanoTag, 1:400), rabbit anti-HA antibody (Alexa Fluor® 555 conjugate, Cell Signaling Technology, # 55420, 1:200), rabbit anti-HA antibody (Cell Signaling Technology, #3724, 1:200). Cells were washed three times with PBS, and then incubated in blocking buffer consisting of 5% bovine serum albumin (Sigma A7906) and 0.2% Triton X-100 (Sigma) in PBS for 1 hour at room temperature with gentle shaking. After permeabilization, cells were incubated with primary antibodies in blocking buffer at 4°C overnight without shaking. Primary antibodies for confocal and TIRF imaging were: chicken anti-MAP2 (Encor, CPCA-MAP2, 1:1,000), guinea pig anti-VGluT1 (Sigma, AB5905, 1:1,000), rabbit anti-Homer1 (Sigma, ABN37, 1:1,000), and FluoTag-X2 anti-PSD95 nanobody (Alexa647 conjugated, NanoTag, 1:500). Primary antibodies for STED imaging of endogenous HA-Lphn3 samples were: mouse anti-Homer1 (Synaptic Systems, #160011, 1:500) and guinea pig anti-VGluT1 (Sigma, AB5905, 1:500). Primary antibodies for STED imaging of lentivirally expressed ALFA-Lphn3 constructs were: rabbit anti-Homer1 (Sigma, ABN37, 1:500) and guinea pig anti-VGluT1 (Sigma, AB5905, 1:500). After three washes with PBS, samples were stained with secondary antibodies in blocking buffer at room temperature (RT) for 1 hour, with gentle shaking. Secondary antibodies used for confocal and TIRF imaging were: anti-chicken AlexaFluor 405, anti-guinea pig AlexaFluor 488, anti-rabbit AlexaFluor 555 (Cell Signaling Technology, all at 1:1000 dilution). Secondary antibodies used for STED imaging of endogenous HA-Lphn3 samples were: anti-rabbit STAR Orange, anti-mouse STAR Red, and anti-guinea pig STAR 460L (Abberior, all at 1:250 dilution). Secondary antibodies for STED imaging of lentivirally expressed ALFA-Lphn3 constructs were: anti-rabbit STAR Orange and anti-guinea pig STAR 460L (Abberior, all at 1:250 dilution). Cells were finally washed 3 times with PBS. For confocal and TIRF samples, cells were kept in glass bottom plates at 4°C in PBS protected from light until imaging (within one month). For STED samples, cells were seeded on #1.5 round coverslips. After staining, cells were briefly washed with distilled water, mounted onto glass slides with ProLong Gold antifade mountant (Thermo Fisher Scientific, #P10144), dried in the dark for 2-3 days, and then sealed with nail polish (Amazon, B000WQ9VNO).

**Confocal, widefield and TIRF imaging**

Unless otherwise noted, all confocal and widefield imaging was conducted using an inverted microscope (Nikon Ti2) equipped with the Celesta 7-line light engine, Crest V3 spinning disk, Kinetix sCMOS camera, and Queensgate Piezo z-drive. Channel settings for different wavelengths are: DAPI (365 nm excitation laser, DAPI/FITC/TRITC/CY5/CY7 multi-band emission filter), GFP (488 nm excitation laser, 511/20 emission filter), RFP (561 nm excitation laser, 595/31 emission filter), and CY5 (640 nm excitation laser, 685/40 emission filter). When imaging widefield fluorescence, the spinning disk was moved out of the light path.

For TIRF imaging, an inverted microscope (Nikon Ti2) equipped with the Ti2-LAPP motorized TIRF module, Kinetix sCMOS camera, and a 60x Apo TIRF (numeric aperture 1.49) were used. Channel settings for different wavelengths were: GFP (488 nm excitation laser, 525/50 emission filter), RFP (532 nm excitation laser, 600/50 emission filter), and CY5 (640nm excitation laser, 685/40 emission filter). To image dendrites that were not immediately adjacent to the glass coverslip, the TIRF illumination angle was reduced to perform highly inclined and laminated optical sheet (HILO) imaging.

When imaging live cells, the microscope environmental chamber was maintained at 37 °C with 5% $CO_2$.

**Single molecule sample preparation and TIRF imaging**

For single-molecule counting of the anti HA-AlexaFluor555 antibody, #1.5 glass coverslips were cleaned by sequentially dipping them into 100% ethanol and Milli-Q water, followed by brief flaming. The coverslips were then coated with BSA for 5 minutes, rinsed with Milli-Q water, and allowed to dry completely before flow-cell chamber assembly. Imaging chambers were constructed by applying double-sided tape to a microscope slide and placing a cleaned coverslip on top to form a flow channel. Antibody samples were prepared in size-exclusion chromatography (SEC) buffer with 3000x dilution of mercaptoethanol (BME). The anti-HA-AlexaFlour555 antibody stock was briefly spun down and diluted 1:50 in the SEC+BME solution, then centrifuged for 5 minutes at 16,000 x g. The diluted antibody solution was loaded into the chamber and allowed to absorb to the coverslip for 30 minutes. Photobleaching of the sample was performed in TIRF with RFP channel settings, with a 300-ms exposure time, 30% laser power, and 2x2 binning.

**Neuron TIRF imaging and single-molecule counting analysis**

Cultured neurons were fixed and stained in 24-well glass bottom plates (Cellvis) and stored in PBS before imaging under TIRF illumination. Rectangular regions were selected from whole fields of view (FOVs) where all channels of the neuron sample (HA-Lphn3-AlexaFluor555, VGluT1-AlexaFluor488, and psd95-AlexaFluor647) were in focus. Single-frame snapshots of all three channels were acquired before acquiring time-lapse photobleaching movies in the RFP channel (HA-labeled exogenous Lphn3). Imaging settings were: 640nm (laser power 50%, exposure time 300ms), 532nm (laser power 80%, exposure time 350ms), and 488nm (laser power 3%, exposure time 200ms), without binning.

Similar analysis pipelines were used to analyze the photobleaching data from the anti HA-AlexaFluor555 single-molecule sample and the stained neuron samples. Rolling-ball background subtraction was applied to all frames in Fiji with a neighborhood size of 25 pixels. Bleaching trace analysis and molecule counting were conducted with custom-written Python scripts using a pipeline similar to that described in previous work[4]. Briefly, each frame went through a 3x3 mean filter, followed by median filter along the time axis with a 5-frame window size. Masks of individual puncta were detected using global thresholding (single molecule samples) and adaptive thresholding (neuron samples) followed by watershed thresholding on the first frame, as x-y drift was negligible throughout the time series. Bleaching traces were calculated by summing the pixel intensities within each punctum mask. To further remove noise, raw traces were passed through a Chung-Kennedy filter to preserve steps[5]. Step sizes of individual fluorophores were determined as follows. Pairwise differences (PPD) between all later and earlier frames were calculated from filtered traces:

$\Delta I_{ij} = I_{ti} - I_{tj}$, for all data pairs where $ti > tj$.

We then calculated the single-sided power spectrum of the PPD distribution using a specified bin size. Automatic peak detection was performed on the power spectrum using scipy.signal.find_peaks. The first peak detected corresponds to the dominant frequency ($f_{dominant}$). The step size for each trace was calculated using $step\ size = {}^{bin\ size}/_{f_{dominant}}$.

The number of fluorescent molecules was estimated as ${}^{(I_{pre-bleach} - I_{post-bleach})}/_{step\ size}$, rounded to the nearest integer, in which $I_{pre-bleach}$ and $I_{post-bleach}$ were calculated from averaging the first and last 5-20 frames. For calculating Lphn3 cluster size in neuron samples, the number of AlexaFluor555 for each cluster was measured as explained above. The number of Lphn3 molecules per cluster was estimated by dividing the number of AlexaFluor555 per cluster (in the neuron sample) by the average number of AlexaFluor555 per HA antibody (measured from the single-molecule antibody sample, Fig. S1F&G).

For stained neuron samples, snapshots from the pre- and post-synaptic marker channels (VGluT1-AlexaFluor488 and psd95 nanobody-AlexaFluor647) were used to identify the locations of excitatory synapses. A scaler background was removed from each channel before applying adaptive thresholding methods to detect pre- and post-synaptic puncta. For each detected Lphn3 punctum, the centroid-to-centroid distance to the nearest pre- or postsynaptic punctum was calculated.

### Glutamate imaging with iGluSnFR3 and data analysis

For glutamate imaging, DIV14 primary neurons were seeded in 24-well glass bottom plates (Cellvis). Endogenous HA-Lphn3 was surface labeled with anti-HA-AlexaFluor555 for 30 minutes. The medium was replaced with warm imaging buffer (Tyrode's solution, Thermofisher J67607.AP, supplemented with 2 mM $CaCl_2$ and 3 mM KCl, final concentration 4 mM $CaCl_2$ and 8 mM KCl) before imaging by TIRF microscopy. Snapshots from the RFP (HA-Lphn3) and GFP (iGluSnFR3) channels were acquired before acquiring the GFP times series. Imaging settings were RFP (80% laser power, 350 ms, no binning), and GFP (10% laser power, 50 ms, 2x2 binning). iGluSnFR3 time series were acquired every 50 ms for 1 minute.

Fiji and custom Python scripts were used for analysis. The iGluSnFR3 channel was first processed with the scale function in Fiji to match the x-y dimensions of that from the HA-Lphn3 channel. Rectangular regions (dendritic segments) were selected from whole FOVs where both channels (HA-Lphn3-AlexaFluor555 and iGluSnFR3) were in focus. To generate Lphn3 cluster masks, adaptive thresholding followed by watershed thresholding was applied to the RFP (Lphn3) channel. The mean fluorescence intensity of each cluster was then extracted as an estimate of cluster size. To generate excitatory synapse masks from the GFP (iGluSnFR3) channel, a maximum-intensity projection over time was created and then synapses were masked with adaptive thresholding. This analysis detects excitatory synapses that have fired at least once during iGluSnFR3 time series imaging. Average intensity traces were extracted from each detected synapse mask over time and then normalized to their respective baselines. iGluSnFR3 spikes were detected using scipy.find_peaks function. To account for variability in spike frequencies in different segments, average Lphn3 cluster intensity and average iGluSnFR3 frequency of each segment were used to calculate normalized Lphn3 cluster intensities and iGluSnFR3 frequencies (Fig. 2B, C).

### Analysis for light-induced Lphn3 cluster formation in live neurons

Analysis was conducted with custom Python scripts on widefield fluorescence time series of live neurons that were surface labeled with ALFA nanobody conjugated with AlexaFluor647 (NanoTag Biotechnologies, N1502). Bleaching correction was performed using Fiji, and a scalar background was subtracted from each image. Then, adaptive thresholding was then used to generate dendrite masks, from which pixel intensities were extracted. Histograms weighted by pixel intensity were calculated for each time frame with 60 bins, followed by one-dimensional Gaussian smoothing, which were then used for plotting (Fig. S2C&D).

### Synapse confocal imaging and synapse density analysis

DIV14 cultured neurons were stained for ALFA-Lphn3 (AlexaFluor647), VGluT1 (AlexaFluor488), Homer1(AlexaFluor555), and MAP2 (AlexaFluor405). Z-stack confocal images were acquired using a 60x oil Plan Apo objective (Nikon) with 0.5 µm z-step for a total of 17 steps. Channels were: CY5 (50% laser power, 250ms exposure), RFP (50% laser power, 250ms exposure), GFP (40% laser power, 150ms exposure), and DAPI (8% or 10% laser power, 100ms exposure).

Maximum-intensity z-projection images were generated with Fiji. Secondary or tertiary dendritic segments were cropped for synapse density analysis. Custom-written Python script was used for automatic image analysis. A scalar background was first subtracted from each channel. Masks were then generated from the Lphn3, VGluT1, and Homer1 channels by adaptive thresholding (Skimage[6]). To ensure consistency, the same segmentation settings were used across conditions in the same experimental batch. Synapses were defined by the overlap of VGluT1 and Homer1 masks, and synapse density was calculated from synapse number divided by dendrite length. All processing steps up to automatic segmentation were performed in a blinded manner.

### 3D STED imaging and analysis

3D-STED microscopy was performed using a Facility Line STED system (Abberior) coupled to an Olympus IX83 microscope through a 60x oil objective (Olympus X-Apo, NA 1.42). Pulsed excitation lasers at 488 nm, 595 nm, and 640 nm were used for fluorophore excitation, and a 775 nm pulsed STED laser was used for depletion. Images were acquired using the 75% 3D-STED setting. The pixel size was set to 40 nm × 40 nm in the xy plane, and z-stacks were collected with a step size of 65 nm. All image acquisitions were performed blinded to experimental conditions to eliminate bias.

Raw STED image stacks of dendritic segments were imported into Imaris (Oxford Instruments) for quantitative analysis. Three-dimensional stacks of Lphn3, VGluT1, and Homer1 channels were smoothed before segmentation. Adaptive thresholding followed by object-splitting was applied to generate 3D surfaces. The same segmentation parameters were used across all images in each experimental batch to ensure consistency. The volumes of the Lphn3, Homer1, and VGluT1 puncta as well as the shortest surface-to-surface distances between the segmented Lphn3 puncta and pre- or post-synaptic marker surfaces were calculated through the software. Puncta diameters were estimated from 3D volumes assuming perfect sphere.

### Calcium imaging and data analysis

DIV14 cultured neurons expressing jRGECO1a were used for calcium imaging. The medium was replaced with warm imaging buffer (Tyrode's solution, Thermofisher J67607.AP, supplemented with 2 mM $CaCl_2$ and 3 mM KCl, final concentration 4 mM $CaCl_2$ and 8 mM KCl) before imaging with widefield microscopy. Time series were acquired at 100 ms interval for 2 minutes in each field, with 4-5 fields per condition. Images were taken with a 20x Air Plan Apo objective (Nikon), and settings were 4x4 binning, RFP (3% laser power, 100 ms exposure).

Fiji and custom Python scripts were used for analysis, similar to a previous method[7]. Maximum time projection was generated for cell masking with the Trackmate-Cellpose plugin in Fiji[8–10]. A scalar background was subtracted, and mean intensity from each cell was extracted overtime and then normalized to its respective baseline level. Population-level calcium traces were calculated by averaging normalized traces from all detected cells in each field-of-view. Spikes from population-level traces were detected using scipy.find_peaks function. Calcium spiking frequency was calculated by the number of spikes divided by the total recording time, and spiking amplitude was calculated as the mean height of detected spikes. We excluded fields-of-view containing less than 20 cells or more than 100 cells to avoid cell densities that were too low or too high. To calculate normalized frequency and amplitude, the average value from all fields of view in the no-stimulation condition within each experiment was used for normalization.

**cAMP signaling assay using G-Flamp1 reporter**

HEK293T cells were seeded in PLL-coated glass bottom 96-well plates (Cellvis P96-1.5H-N) at 50-60k cells/cm$^2$ and allowed to adhere overnight. The next day, 30 ng reporter plasmid (ePBCAG-bsd-mScarlet-I-G-Flamp1) and 40 ng Lphn3 (ePBCAG-IgK-HA-ALFA-hsLphn3-mIRFP670/ ePBCAG-IgK-HA-ALFA-hsLphn3-F1041E-mIRFP670) or vector plasmids were transfected in each well using Lipofectamine 3000 reagents (Invitrogen). 24-27 hours post transfection, medium was changed to imaging media (DMEM with HEPES, no phenol red, Gibco 21063029). Cells were kept in imaging media for 30 minutes to 1 hour before being transferred to a 37°C microscope for imaging assays.

An inverted microscope (Nikon Ti-E) equipped with the Heliophore (89 North) LED engine and Andor Neo sCMOS camera was used to conduct widefield fluorescence imaging. Images were acquired through the Nikon 20x air PlanApo objective. Images in GFP and RFP channels were acquired every 5 minutes for a total of 6 frames, with 50 μM forskolin (MilliporeSigma 34-428-25MG) added after the second frame to confirm proper reporter function.

Custom-written Python script was used to analyze data. Single-cell segmentation and tracking were performed using the Trackmate-Cellpose plugin in Fiji[8–10] using the RFP (mScarlet-I) channel. Scalar background values were subtracted from each channel before we calculated the normalized reporter image through pixel-by-pixel division of the GFP (G-Flamp1 reporter) and RFP (expression normalization) channels. To prevent artifacts resulting from small values in certain denominator pixels from the RFP channel, pixel values below 1 were set as 1. Mean intensities from each cell were extracted from the normalized reporter image as the cAMP reporter signal. To validate the G-Flamp1 reporter, we added 50μM forskolin and observed that the GFP signal went up and plateaued within 15 minutes (Fig. S4 A, B).

**TRUPATH assay**

HEK 293T cells were plated in poly-L-lysine-coated, white, opaque, Nunclon Delta-Treated Flat-Bottom 96-well assay plates (Thermo Fischer Scientific) at a density of 30-50k cells/cm$^2$. After 18-24 hours, cells in each well were transfected with 200 ng of receptor DNA (HA-Lphn3-mIRFP670, WT or F1041E) and 100 ng DNA of trimeric G-protein mix containing Gα-RLuc8:Gβ:Gγ-GFP2 at 1:1:1 mass ratio. Gα-RLuc corresponded to Gαs, Gα12, or Gα13. In vehicle control wells, an empty vector was used for transfection. Transfection was conducted using the calcium phosphate method, in which a HEPES-buffered saline solution was used to facilitate precipitation of DNA-calcium phosphate complexes, which were then added dropwise to the cells. 26-30 hours after transfection, the cells were serum-starved by replacing the serum-containing HEK growth media with 65 uL of warm FluoroBrite DMEM buffer (Gibco A1896701) in each well. Cells were incubated for 1 hour followed by addition of 10 µl of freshly diluted 100 µM Prolume Purple (methoxy e-Coelenterazine; Nanolight Technologies), which was stored as 1mM stock in methanol at -80 °C.

After a 5-7 min equilibration period, plates were read in a BioTek Synergy plate reader using 460/40 nm (RLuc8) and 535/35 nm (GFP2) emission filters, at gain settings of 160 and 200, respectively, with signal integration time of 2.5 s per well. Plates were read row by row, from 7 minutes to 20 minutes after Prolume Purple addition. Measurements collected between 10 and 15 min were used for analyses. Each condition was tested in triplicate per plate, and in each replicate, Prolume purple was added 5-7 min before signal readout.

BRET2 ratios were calculated as the mean ratio of the GFP2 emission to RLuc8 emission. BRET values from each Gα condition were normalized to their respective vector controls of the same plate-reading run to quantify relative changes in signaling.

**Surface labelling of Lphn3 receptor in HEK cells and surface expression measurement**

Surface labeling of the transfected HA-Lphn3 receptor constructs was conducted in parallel to the TRUPATH assay with the same batches of HEK293T cells, except cells were plated in poly-L-lysine-coated, glass-bottom 96-well assay plates (Cellvis). 26 to 30 hours after transfection, HEK growth media from each well was carefully pipetted out and replaced with 50uL of warm FluoroBrite DMEM buffer (Gibco) containing anti-HA antibody conjugated to Alexa-fluor-555 (1:200 dilution, Cell Signaling Technology, 55420). Cells were incubated at 37°C for 15-20 min. Cells were then fixed by 4% paraformaldehyde (PFA), prepared freshly from 16% stock in 1x PBS (pH = 7.0), incubated at room temperature for 10 minutes, and washed twice with FluoroBrite DMEM. Finally, cells were imaged at room temperature on the spinning disk confocal microscope using 20x or 40x Nikon PlanApo air objectives in the RFP channel.

Custom-written Python script was used to analyze data. A scalar background was first subtracted. Masks were then generated with adaptive thresholding (scikit-image) to extract mean intensities. For each transfection batch, surface receptor intensities were normalized to the average signal from HA-Lphn3-WT before plotting the results.

**Halo-AKAR PKA reporter assay and analysis**

NIH3T3 mouse fibroblasts were chosen for this assay over HEK293T cells due to their low expression of exogenous genes upon transient transfection and their flat morphology. These properties allow us to achieve moderate levels of basal receptor clustering and to monitor receptor clustering on the cell surface.

Fibroblast cells were seeded in PLL-coated glass bottom 96- well plates (Cellvis P96-1.5H-N) at 20-30k cells/cm$^2$ and allowed to attach overnight. The next day, 120ng reporter plasmid (Halo-AKAR2.1) and 150ng Lphn3/vector plasmids were transfected in each well using Lipofectamine 3000 reagent (Invitrogen). 23-25 hours post transfection, Janelia Fluor 635 HaloTag Ligand (gift from Dr. Luke Lavis, Janelia Research Campus) was added to medium at 500 nM and cells were incubated for 2-3 hours. Then, medium was changed to imaging solution (DMEM with HEPES, no phenol red, Gibco 21063029). Cells were kept in imaging solution for 30 minutes to 1 hour before transferred to a microscope for imaging assays.

Confocal images in CY5 and RFP channels were taken with the 60x oil PlanApo objective (Nikon) with 2x2 binning. Imaging settings were: CY5 (70% laser power, 300 ms exposure) and RFP (60% laser power, 200 ms exposure). Time series was taken at 20 second interval. Blue light stimulation was applied in one pulse with 50% power with the 488 nm laser for 2 seconds. To induce maximal reporter signal, a mixture of IBMX (Tocris 28-455-0) and forskolin (MilliporeSigma 34-428-25MG) at final concentrations of 100 μM and 50 μM respectively, was added towards the end of imaging.

Custom-written Python script was used to conduct image analysis. First, a scalar background was subtracted from each channel followed by whole cell mask generation with adaptive thresholding (scikit-image) to extract mean reporter intensity for each frame. A ring mask with thickness of 4 pixels (0.8668 μm) was generated from the cell mask to extract membrane Lphn3 intensity. Reporter signal was normalized by ${}^{\Delta F}/_{\Delta F_{max}}$. To ensure analysis of cells with functional reporter activity, only traces with normalized reporter value at the last frame exceeding 0.5 were retained for analysis.

### References (for Methods)


1. Sando, R., Jiang, X. & Südhof, T. C. Latrophilin GPCRs direct synapse specificity by coincident binding of FLRTs and teneurins. *Science* **363**, eaav7969 (2019).
2. statannot: add statistical annotations on an existing boxplot/barplot generated by seaborn.
3. Wang, S. *et al.* Alternative splicing of latrophilin-3 controls synapse formation. *Nature* **626**, 128–135 (2024).
4. Nissen, S. B. *et al.* Cluster assembly dynamics drive fidelity of planar cell polarity polarization. *Nat Commun* https://doi.org/10.1038/s41467-026-71839-5 (2026) doi:10.1038/s41467-026-71839-5.
5. Chung, S. H. & Kennedy, R. A. Forward-backward non-linear filtering technique for extracting small biological signals from noise. *Journal of Neuroscience Methods* **40**, 71–86 (1991).
6. Walt, S. van der *et al.* scikit-image: image processing in Python. *PeerJ* **2**, e453 (2014).
7. Sun, Z. & Südhof, T. C. A simple Ca2+-imaging approach to neural network analyses in cultured neurons. *Journal of Neuroscience Methods* **349**, 109041 (2021).
8. Stringer, C. & Pachitariu, M. Cellpose 2.0: how to train your own model. 2022.04.01.486764 Preprint at https://doi.org/10.1101/2022.04.01.486764 (2022).
9. Tinevez, J.-Y. *et al.* TrackMate: An open and extensible platform for single-particle tracking. *Methods* **115**, 80–90 (2017).
10. Zhu, X., Smith, M. V. E. & Tanentzapf, G. Automated detection of spontaneous calcium signaling events in prohemocytes of the Drosophila melanogaster lymph gland. *MicroPubl Biol* **2025**, 10.17912/micropub.biology.001883.
